# Structure and mechanism of the type I-G CRISPR effector

**DOI:** 10.1101/2022.08.08.503147

**Authors:** Qilin Shangguan, Shirley Graham, Ramasubramanian Sundaramoorthy, Malcolm F White

## Abstract

Type I CRISPR systems are the most common CRISPR type found in bacteria. They use a multisubunit effector, guided by crRNA, to detect and bind dsDNA targets, forming an R-loop and recruiting the Cas3 enzyme to facilitate target DNA destruction, thus providing immunity against mobile genetic elements. Subtypes have been classified into families A-G, with type I-G being the least well understood. Here, we report the composition, structure and function of the type I-G Cascade CRISPR effector from *Thioalkalivibrio sulfidiphilus*, revealing key new molecular details. The unique Csb2 subunit processes pre-crRNA, remaining bound to the 3’ end of the mature crRNA, and seven Cas7 subunits form the backbone of the effector. Cas3 associates stably with the effector complex via the Cas8g subunit and is important for target DNA recognition. Structural analysis by cryo-Electron Microscopy reveals a strikingly curved backbone conformation with Cas8g spanning the belly of the structure. Type I-G Cascade is one of the most streamlined Class 1 CRISPR effectors. These biochemical and structural insights shed new light on the diversity of type I systems and open the way to applications in genome engineering.

## INTRODUCTION

Type I CRISPR effectors, which have the signature protein Cas3, are the most abundant CRISPR systems found in prokaryotes (1). The canonical features include a Cas7 backbone that binds crRNA, a large subunit (Cas8) that is involved in detection of the protospacer adjacent motif (PAM) in DNA targets, a Cas6-like nuclease that processes crRNA and remains bound to the 3’ end, a Cas5 subunit in single copy at the 5’ end of the crRNA and a small subunit (Cas11) (2). Cas5, 6 and 7 are all members of the “RRM” (RNA Recognition Motif) superfamily (3). On binding target DNA with a suitable PAM and forming an R-loop, the type I effector (also known as “Cascade” (4)) recruits the Cas3 protein, which loads on to the non-target strand and translocates with a 3’ to 5’ polarity, degrading the DNA with its HD-nuclease domain (5,6).

High resolution crystal or cryo-EM structures are available for type I-A (7), I-C (8,9), I-D (10), I-E (11–14), I-F (15), I-Fv (16) and type I-F-TniQ (17) complexes. However, within this broad family there are many variations. The type I-B, I-C and I-D systems encode the small Cas11 subunit within the gene for the large subunit (18) while the type I-F and I-G systems lack a small subunit altogether (15). The type I-D system represents a half-way house between type I and III systems, with a large subunit related to Cas10 rather than Cas8 (19,20). In type I-C systems, the Cas5 subunit possesses crRNA processing activity and Cas6 is absent (21). Although more complex than the streamlined class 2 systems such as Cas9 and Cas12, type I effectors have been utilised for a growing range of genome editing approaches and are particularly useful for long range genome edits (7,22–25).

The only type I effector that has not been characterised on a biochemical or structural level is type I-G (previously known as type I-U) (26). Type I-G is a minimal system lacking a *cas11* gene and with a predicted fusion of Cas5 and Cas6 in a subunit known as Csb2 (1). This system has been shown to use a TTN PAM to provide interference against targeted plasmids *in vivo* (27). The large subunit, known as Csx17, Cas8u (26) or hereafter Cas8g is highly diverged from any of the structurally characterised large subunits of type I and III systems. Here, we show that the type I-G system from *Thioalkalivibrio sulfidiphilus* HL-EbGr7 (Tsu) assembles into a functional Cascade complex that provides efficient interference against target plasmids or phage in conjunction with Cas3, which is an integral subunit of the effector. The C-terminal domain of the Csb2 subunit is shown to function as the crRNA processing endoribonuclease, and we see no evidence for a small subunit encoded within the large subunit Cas8g. The cryoEM structure of type I-G Cascade reveals highly curved backbone organisation most reminiscent of type I-F effectors, and a unique subunit organisation.

## MATERIAL AND METHODS

### Cloning

Synthetic genes encoding Cas proteins (Cas8g, Csb2, Cas7 and Cas3) were obtained from Integrated DNA Technologies (Coralville, IA, USA). Restriction enzyme sites were added when necessary and codon usage was optimized for *Escherichia coli. Cas8g, cas7* and *cas3* genes were digested with *Nco*I and *Bam*HI (Thermo Scientific) and ligated into pEV5HisTEV (28) to produce the vector that allows expression of individual proteins with N-terminal TEV cleavable His_8_-tags. *csb2* was cloned into the pET-Duet (Novagen, Merck Millipore) vector via ligation after *Nde*I and *Xho*I (Thermo Scientific) digestion. Site directed mutagenesis of *cas* genes was carried by standard protocols using Phusion enzyme (Thermo Scientific).

A CRISPR array containing 6 identical spacers targeting the *tetR* gene flanked by 7 repeats was cloned into pCDF-Duet (Novagen, Merck Millipore) by ligation after *Nco*I and *Sal*I digestion. The other two CRISPR arrays: *lacZ* target (5 repeat, 4 spacer) and *lpa* target (4 repeat, 3 spacer) were prepared using the same method. Supplementary Table 1 shows the sequences.

To express the type I-G complex for *in vivo* studies, vector pACE-M1 (MultiColi™, Geneva Biotech, Genève, CH) was assembled by SLIC (sequence and ligation independent cloning). DNA fragments encoding *cas8g, csb2* and *cas7* were amplified with PCR prior to SLIC and ligation into pACE, placing these three genes under control of a single T7 promoter. The *cas*3 gene was digested with *Nco*I and *Sal*I and incorporated into vector pRAT under the control of the araBAD promoter. Plasmid lacZ-pRAT was described previously (29). All final constructs were verified by sequencing (GATC Biotech, Eurofins Genomics, DE)

### Protein expression and purification

pEV5HisTEV vectors harbouring *cas* genes were transformed into *E. coli* C43(DE3) for protein expression. Cells grew in LB culture containing 50 μg ml^-1^ kanamycin overnight. Following a 100-fold dilution, cells were grown at 37 °C, 180 rpm to reach an OD_600_ of 0.6~0.8, induced by 400 μM IPTG, followed by overnight protein expression at 25 °C. Cell pellets were harvested by centrifugation and resuspended in lysis buffer (50 mM Tris-HCl, 250 mM NaCl, 10 mM imidazole, and 10 % glycerol, pH 7.5). The cells were then sonicated, and debris was removed by centrifugation. Cell lysate was loaded to a pre-equilibrated HisTrap Crude FF (GE Healthcare) column, washed with wash buffer (50 mM Tris-HCl, 250 mM NaCl, 30 mM imidazole, and 10 % glycerol, pH 7.5), eluted with elution buffer (50 mM Tris-HCl, 250 mM NaCl, 500 mM imidazole, and 10 % glycerol, pH 7.5). Fractions containing Cas protein were pooled, followed by TEV cleavage to remove the His-tag and passage through the HisTrap column for the second time. The cleaved protein was collected and further purified using gel filtration (HiLoad 16/60 Superdex pg 200, GE Healthcare) with GF buffer (20 mM Tris-HCl, 250 mM NaCl, pH 7.5). Pure Cas protein was finally concentrated in Amicon Ultra centrifugal filter (Merck-Millipore).

### Effector complex reconstruction

Effector complexes were assembled by mixing individual pure protein subunits with pre-crRNA, obtained by *in vitro* transcription (MEGAscript™ T7 Transcription Kit, Invitrogen™) in the combinations noted in the results. The Cas protein combinations were incubated with pre-crRNA in 20 mM Tris-HCl, 250 mM NaCl, 1 mM DDT, 1 mM EDTA and 0.5 U/μl RNase inhibitor pH 7.5 for 1 h at 37 °C, filtered by centrifugation at 10000 x g for 10 min, then loaded onto a Superose 6 10/300 increase (GE Healthcare) column for gel filtration in GF buffer buffer (20 mM Tris-HCl, 250 mM NaCl, pH 7.5). Fractions containing the complex were pooled and concentrated using a centrifugal filter (Vivaspin^®^ 500, MW cutoff 30,000 Dalton, Vivaproducts).

### Csb2 N-terminal domain and C-terminal domain expression

A stop codon was introduced by mutagenesis at R260 of Csb2 to allow the expression of the N-terminal domain on pEV5HisTEV vector. For the C-terminal domain, primers with NcoI and BamHI were used to amplify C-terminal sequence fragment encoding from R262 to the end of the *csb2* gene, which was cloned into the pEV5HisTEV vector by digestion and ligation. Both N- and C-terminal domains were expressed and purified as described for the full-length proteins above.

### Oligonucleotides

All 6-FAM™-labelled and non-labelled DNA or RNA substrates were purchased from Integrated DNA Technologies (Leuven, BE). Where required, oligonucleotides were 5’-end-labelled with [γ-^32^P]-ATP (10 mCi ml^-1^, 3000 Ci mmol^-1^, Perkin Elmer) with polynucleotide kinase (Thermo Scientific). Duplex DNA was obtained by annealing equimolar amount of complementary ssDNA in 10mM Tris-HCl, 50mM NaCl, pH7.5, 95 °C for 5min, slowly cooling down overnight to room temperature in a heat block. All oligonucleotide sequences can be found in Supplementary Table 1.

### CRISPR repeat cleavage assay

36 nt 5’-6-FAM™-labelled CRISPR repeat RNA was incubated with Csb2 or other Cas protein in 20 mM Tris-HCl, 50 mM NaCl, 1 mM DTT, 5 mM EDTA, 0.1 U/μl RNase inhibitor, pH 7.5 for 5 min at 37 °C, at a final concentration of 50 nM RNA and 0.5 μM protein. Reaction was stopped by adding formamide and heat denaturing at 95 °C for 3 min. Product was loaded to a denaturing gel (20% acrylamide, 7M Urea), running in 1 X TBE buffer, and visualized by scanning (Typhoon FLA 7000, GE Healthcare). RNA ladders were obtained by alkaline hydrolysis of the CRISPR repeat RNA (Thermo Fisher Scientific, RNA Protocols).

### pre-crRNA cleavage assay

643nt pre-crRNA was *in vitro* transcribed with [a-^32^P]-ATP (10 mCi ml^-1^, 3000 Ci mmol^-1^, Perkin Elmer) as described above, followed by phenol-chloroform extraction and ethanol precipitation. Reactions contained 1 μM pre-crRNA in reaction buffer (20 mM Tris-HCl, 50 mM NaCl, 1 mM DTT, 0.1 mg/ml BSA, 5 mM MgCl_2_, 1 mM EDTA, 0.1 U/μl RNase inhibitor, pH 7.5). Wildtype or variant H503A Csb2 was added to the reaction at 0.5 μM followed by incubation at 37 °C for 30 min. Cleavage products were resolved in a denaturing gel as described above.

### Fluorescence anisotropy

The method is adapted from (30). In detail, 25 nM 5’-6-FAM™-labelled CRISPR hairpin or repeat RNA was suspended in a quartz cuvette with 50 mM NaCl, 1 mM DTT, 5 mM MgCl_2_, 1 mM EDTA, 0.1 U/μl RNase inhibitor (Thermo Scientific). The initial anisotropy of the RNA was measured in a Varian Cary Eclipse fluorescence spectrophotometer (Agilent Technologies), using the Eclipse ADL application. The measurement was carried at 37 °C, exciting the fluorescein-labelled RNA at 480 nm and monitoring emitted fluorescence at 525 nm. Csb2 was titrated at progressively higher concentrations into the sample. All points of titration were carried out with automatic polarizers. The anisotropy value of each point was plotted against Csb2 concentration, and a curve was fitted to get *K*_D_ value using equation below.

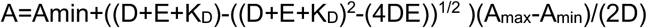

Where A is anisotropy of free RNA, D is total RNA concentration, E is total protein concentration. A_max_ and A_min_ is maximum and minimum anisotropy. The equation assumes a RNA:protein binding stoichiometry of 1:1 (30).

### Electrophoretic Mobility Shift Assay (EMSA) of short dsDNA

10 nM [γ-^32^P]-labelled dsDNA was mixed with 0.8 μM effector complex in 10 μl reaction buffer (20 mM Tris-HCl, 50 mM NaCl, 1 mM DTT, 0.1 mg/ml BSA, 5 mM MgCl_2_, 1 mM EDTA) at 37 °C for 30 min, loaded onto an 8 % acrylamide 1 x TBE gel with Ficoll loading buffer, and electrophoresed at 200 V for 1 h, followed by visualization (Typhoon FLA 7000, GE Healthcare).

### Plasmid DNA binding and cleavage assays

*In vitro* reconstructed effector complex was mixed with target or control plasmid (2 nM), incubated in 10 μl reaction buffer (20 mM Tris-HCl, 50 mM NaCl, 1 mM DTT, 0.1 mg/ml BSA, 5 mM MgCl_2_, 1 mM EDTA) at 37 °C for 1 h. ATP was present at 2 mM where indicated. Reactions were analysed by separation on a 0.8 % Agarose gel, running in 1 X TBE buffer at 10 mA overnight, post-stained with SYBR green for 30 min, and visualized by scanning (Typhoon FLA 7000, GE Healthcare). Open circular plasmid control was obtained by incubation with the *Nt.BspQI* nickase (New England Biolabs), and linear plasmid by cleavage with *Bam*HI.

### Analysis of dsDNA cleavage

*In vitro* reconstructed effector complex was mixed with 16.6 nM [γ-^32^P]-labelled dsDNA and incubated in reaction buffer (20 mM Tris-HCl, 50 mM NaCl, 1 mM DTT, 0.1 mg/ml BSA, 5 mM MgCl_2_, 1 mM EDTA) at 37 °C for 1 h, supplemented with 2 mM ATP as indicated. Reactions were stopped by addition of 0.5 M EDTA, an equal volume of formamide and denaturation at 95 °C for 3 min. Cleavage products were separated on a S2 sequencing gel (20 % acrylamide, 7 M Urea), 90 W, 70 min and visualized by scanning as above. A Maxam-Gilbert G+A ladder was acquired by incubating 5 ng ^32^P-labelled oligonucleotide with 1 μg calf thymus DNA and 0.4 % formic acid in 10 μl TE buffer (10 mM Tris-HCl, 0.1 mM EDTA, pH 7.5) at 37 °C for 25 min, followed by addition of 150 μl 1 M piperidine and heating at 95 °C for 30 min. The ladder was ethanol precipitated before resuspension into loading buffer and loading onto the gel.

### Plasmid challenge assay

The method was described previously (29). For the type I-G system, pACE-M1 and pCDF with targeting CRISPR array were co-transformed into *E. coli* C43 (DE3). Transformants were selected by 100 μg ml^-1^ ampicillin and 50 μg ml^-1^ spectinomycin. Competent cells were prepared by diluting an overnight culture 50-fold into fresh, selective LB medium. The culture was incubated at 37 °C, 220 rpm to reach OD_600_ 0.4 to 0.5. Cells were collected by centrifugation and the pellet resuspended in an equal volume of 60 mM CaCl_2_, 25 mM MES, pH 5.8, 5 mM MgCl_2_, 5 mM MnCl_2_. Following incubation on ice for 1 h, cells were collected and resuspended in 0.1 volumes of the same buffer containing 10 % glycerol. Aliquots were stored at −80 °C. pRAT Plasmid with or without Cas3 was transformed to the competent cells. Transformation mixture with LB medium was incubated with shaking for 2.5 h after heat shock. A total of 3 μl transformation product was applied in a 10-fold serial dilution to LB agar plates supplemented with 100 μg ml^-1^ ampicillin and 50 μg ml^-1^ spectinomycin when selecting for recipients only; transformants were selected on LB agar containing 100 μg ml^-1^ ampicillin, 50 μg ml^-1^ spectinomycin, 25 μg ml^-1^ tetracycline. LB agar plates containing 0.2 % (w/v) D-lactose and 0.2 % (w/v) L-arabinose were used for induction. Plates were incubated at 37 °C for 16–18 h. Colonies were counted manually and corrected for dilution and volume to obtain colony-forming units (cfu) ml^-1^; statistical analysis was performed with Prism8 (GraphPad). The experiment was performed with two biological replicates and at least two technical replicates.

### Phage immunity assay

The method was described previously (31). For the type I-G system, pACE-M1, pCDF-Lpa (CRISPR array targeting phage P1 *Lpa)* and pRAT-Cas3 were co-transformed to *E. coli* C43 (DE3). Cells were selected by 100 μg ml^-1^ ampicillin, 50 μg ml^-1^ spectinomycin and 12.5 μg ml^-1^ tetracycline. The cells were grown overnight at 37 °C in LB medium containing 50 μg ml^-1^ ampicillin, 25 μg ml^-1^ spectinomycin and 12.5 μg ml^-1^ tetracycline. The overnight culture was diluted to OD_600_ of 0.1 by LB medium supplemented with the antibiotics, 10 mM MgSO_4_, 0.2 % (w/v) D-lactose and 0.2 % (w/v) L-arabinose. In uninduced tests, D-lactose and L-arabinose were absent. 160 μl of diluted culture was infected with 40 μl diluted bacteriophage P1 to give MOIs of 1, 0.1 and 0.01 in a 96-well plate. The OD_600_ of the culture in the plate was measured by a FilterMax F5 Multi-Mode Microplate Reader (Molecular Devices) every 20 min over 20 h. The experiment was carried out with two biological replicates and three technical replicates. The OD_600_ was plotted against time using Graphpad Prism 8.

### CryoEM sample preparation and data collection

Pre-formed Type IG complex containing Cas7, Cas8g, Csb2 and crRNA sample was diluted to 1 mg/ml and vitrified in liquid ethane using Vitrobot Mark IV (Thermo Fisher Scientific). Holey carbon Quantifoil (R2×2) grids (EMS) were glow discharged in air using a Quorum technology instrument for 60 seconds at atmospheric pressure of 0.1 prior to application of the sample. 4 μL of the sample applied to the grids at a temperature of 4 °C and a humidity level of 100 %. Grids were then immediately blotted (force 3.5, time 3 seconds) and plunge frozen into liquid ethane cooled to liquid nitrogen temperature. Grids were pre-screened in the in-house JEOL 2200 FS microscope before transported to Electron Bio-imaging centre (eBIC) at the Diamond Light source. Grids were imaged using a 300Titan Krios transmission electron microscope (Thermo Fisher Scientific) equipped with K3 camera (Gatan) operated in super-resolution mode. Movies were collected at 105,000x magnification and binned by two on the camera (calibrated pixel size of 0.84 Å/pixel). Images were taken over a defocus range of −1.5 μm to −3.00 μm with a total accumulated dose of 50e^-^/Å^2^ using EPU (Thermo Fisher Scientific, version-2.11.1.11) automated data software. A summary of imaging conditions is presented in Supplementary Table 2.

### CryoEM data processing

A total of 15710 micrographs were collected and pre-processed using RELION (v3.1) software suite (32)Motion correction of movies was carried out using MotionCor2 (33) and CTF correction was carried out using CTFfind4.1 (34–37) and the classes that contain similar structural features were merged. In the final round of refinement, a total of 415,000 particles has been used. Ctf parameter refinement and Bayesian polishing of the particles were carried out before the very last round of 3D refinement. A detailed workflow of iterative classification and refinement of the particles are shown in Supplementary Figure S6. The seven cas7 subunits with crRNA, which form the crescent structure, were the better-defined component of the complex. The large subunit Cas8g had poorly defined density in the global refinement. To generate a better density of the large subunit we utilised multi-body refinement incorporated in the RELION suite. For this, the Cas7-crRNA was treated as a separate body from the large subunit Cas8g. A separate soft mask of 5 pixels that covered Cas8g and the Cas7-crRNA modules was generated to use it in the multi-body refinement. Multi-body refinement was continued from the last iteration of the consensus refinement, with a sigma angle of 10 and a sigma offset of 2. Principal component analysis for all the particles was performed using RELION-4. Principal components 1 and 2 accounted for 25 % of the variance in the rotations and translations. Post-processing was performed with a soft-mask of 5 pixels and the B-factor estimated automatically in RELION-4 following standard practice. The final resolution of the reconstructed volume was estimated with an FSC correlation at a cutoff of 0.143. The final maps were used for structure fitting and model refinement.

### Model building and refinement

An initial model of each component of the Type I-G complex was generated using Alphafold2 (AF2) (38). Seven of the Cas7 AF2 models were first docked into the density as rigid bodies on a locally sharpened map. The map covering each of the Cas7 rigid body was segmented and the fitting of the model was then continued in the segmented map. Rigid body fitting and segmentation were performed in ChimeraX (37). Subsequently, the initial rigid body of each Cas7 molecule was flexibly refined into the segmented CryoEM map using ISOLDE (v.1.3) (39) implemented in ChimeraX (v.1.3). During refinement we applied adaptive distance restraints to each subunit. Once each Cas7 was individually refined, the Cas7 complex was generated and additional around of ISOLDE refinement was carried out on a non-segmented map to fine-tune the interaction boundaries of each Cas7 molecule and their neighbours. The crRNA was clearly defined in the density. The structure of the crRNA was *de novo* traced in the density using COOT (40). The crRNA structure was then refined in ISOLDE. For the large subunit Cas8g, we used the RELION Multibody refined map for the model fitting. Again, we used the AF2 predicted structure as the starting model. The AF2 model is segmented into three separate regions, the N-terminus (1-381), the middle or bridge region (382-479), and the C-terminus (480-720). These regions were fitted as rigid bodies and then subsequently refined in ISOLDE. We used adaptive secondary structure restraints on the separated regions. The resultant atomic models were then refined using phenix.real_space_refine in Phenix (v.1.20.1-4487) (41) with secondary structure, reference model, and geometry restraints. Model FSC validation tools and the overall quality of the model was assessed in the map using the CryoEM validation tools in Phenix and MolProbity (42). The final refined structures are submitted to the Protein and Electron Microscopy Data Banks with accession numbers PDB 8ANE, 8ANZ and EMDB-15540.

## RESULTS

### Type I-G effector subunit expression and location of pre-crRNA processing activity

*Thioalkalivibrio sulfidiphilus* HL-EbGr7 (Tsu) (43) encodes two CRISPR systems, a type III-B locus array and a type I-G CRISPR locus (Figure 1A). To investigate the uncharacterised type I-G effector, we designed synthetic genes for the expression of each subunit (Csb2, Cas7, Cas8g and Cas3) in *E. coli*, and cloned them in expression plasmid pEV5hisTEV (28), which adds a TEV cleavable polyhistidine tag at the N terminus. The proteins were purified individually using IMAC and size exclusion steps to achieve homogeneity. As the mechanism of pre-crRNA processing in type I-G systems is not known, we incubated Csb2, Cas7 and Cas8g individually with a synthetic RNA containing the 36 nucleotide (nt) CRISPR repeat sequence, observing that Csb2 was the only subunit that cleaved this substrate, generating a mature crRNA with a canonical 8 nt 5’ handle (Figure 1B & 1D). The cleavage site was positioned 4 nt 3’ of the base of the long hairpin formed by the pre-crRNA, an unusual feature as most Cas6 enzymes cleave at the base (44,45). To identify the position of the active site of Csb2, we targeted two conserved histidine residues, replacing them with alanine. The variant proteins were purified as for the wild-type enzyme. The variant containing a H503A mutation no longer cleaved the CRISPR RNA (Figure 1C), pinpointing the active site to the C-terminus of Csb2. This domain has been suggested as a distant homologue of Cas6 by bioinformatic analysis (26). To investigate pre-crRNA processing in more detail, we generated a radioactively labelled pre-crRNA of 643 nt by *in vitro* transcription. Incubation of this transcript with Csb2 generated a series of processed products with a final size consistent with the expected 72 nt of mature crRNAs (Figure S1).

**Figure 1.**
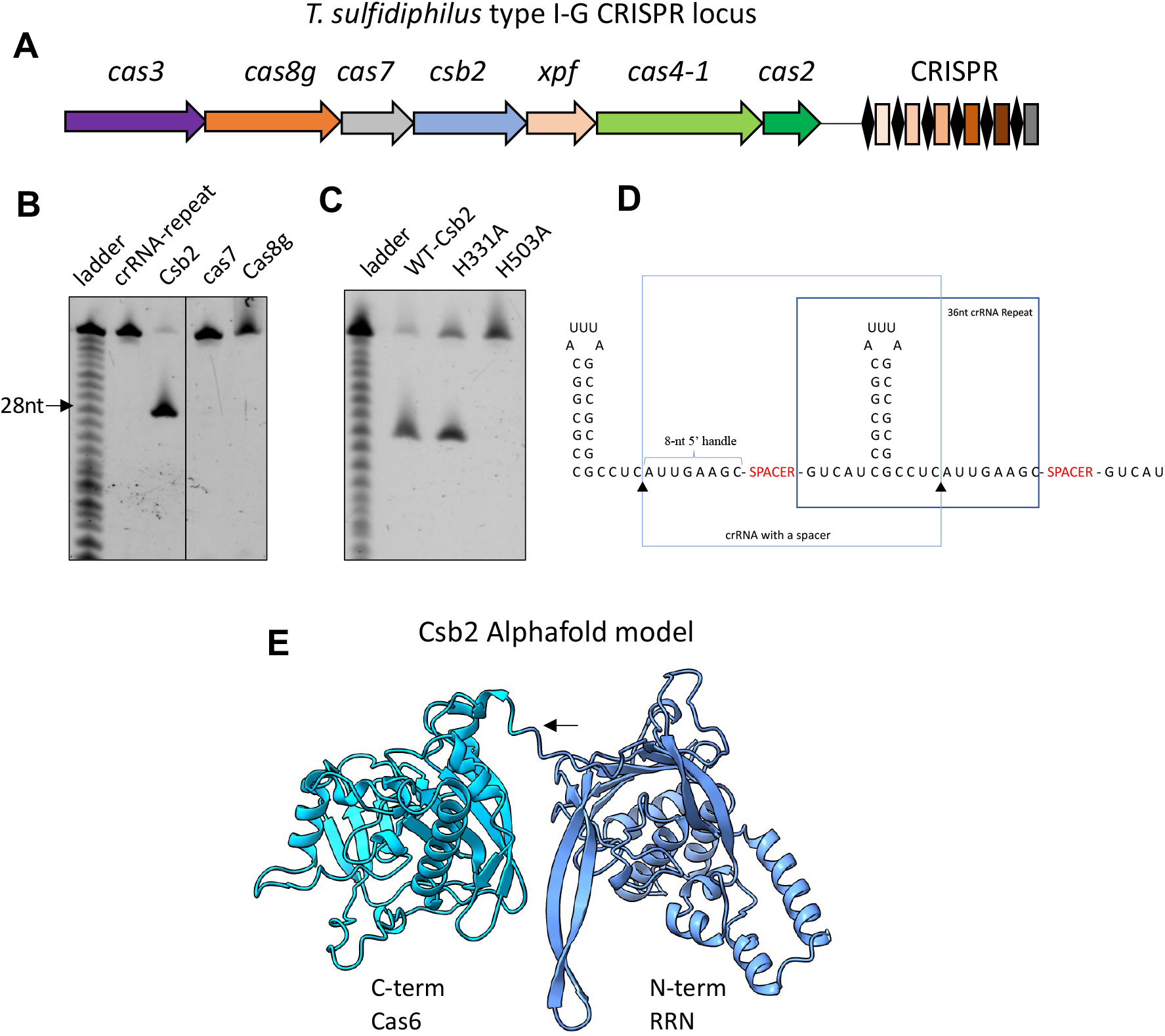
Csb2 generates mature crRNA in type I-G systems. **(A)** An overview of type I-G CRISPR genome locus in *Thioalkalivibrio sulfidiphilus* HL-EbGr7. The *xpf* gene is uncharacterised but most likely encodes a 3’-flap DNA endonuclease involved in prespacer processing for adaptation. **(B)** A synthetic RNA species corresponding to the 36 nt CRISPR repeat was incubated with purified recombinant Csb2, Cas7 and Cas8g at 37 °C for 10 min and analysed by denaturing gel electrophoresis. Only Csb2 generated a cleavage product. The ladder was obtained by alkaline hydrolysis of the crRNA. **(C)** The H503A variant of Csb2 does not cleave the repeat. **(D)** Schematic showing the pre-crRNA with Csb2 cleavage sites indicated. **(E)** Structural model of the Csb2 protein by AF2 suggests a two-domain structure with a C-terminal Cas6-like domain, joined by a linker sequence (Indicated by arrow) to the N-terminal Cas5-like domain. The relative orientation of the two domains cannot be predicted.

To understand the structure of Csb2 in more detail, we used Alphafold2 (AF2) (38) on the Colabfold server (46) to predict the protein structure. This predicted a two-domain structure (Figure 1E). Submitting the model to Dali (47) to look for structural homologues, we observed a good match for the first 150 residues of the N-terminal domain to the Cas5 subunit of type I-E and type III-A CRISPR systems, which has an RRM fold and extended beta-hairpin, and a strong match for the C-terminal domain (Dali Z-score 12.4) to the Cas6b protein (PDB 4Z7K) (48). By overlaying the structure of Cas6b with the AF2 model of the Csb2 C-terminal domain, we could estimate the binding site of the crRNA hairpin, which is consistent with the position of the catalytic H503 residue (Figure S1C).

Although the relative orientation of the two domains of Csb2 cannot be accurately predicted, this gives us a reasonable working model of the Csb2 structure, and provisionally places it at the 3’ end of the crRNA, bound to the extended RNA hairpin of the CRISPR repeat. However, the Cas5d protein of type I-C systems cleaves the CRISPR RNA but remains associated with the 5’ handle rather than the 3’ hairpin, which is subsequently degraded on effector assembly (8). To resolve this uncertainty, we investigated the binding affinity of Csb2 for the 3’ hairpin and the 5’ handle using fluorescence anisotropy. For Csb2, we observed a high binding affinity (K_D_ = 55 nM) for CRISPR RNA (Figure 2A) and a very similar binding affinity (K_D_ = 43 nM) for the 3’ hairpin that is the product of Csb2 cleavage (Figure 2B). By contrast, Csb2 does not bind appreciably to the 8 nt 5’ handle (Figure 2C & S2A). We proceeded to clone and express the separate N- and C-terminal domains of Csb2, using the domain boundary at residue R260 in the linker (Figure 1E). The two domains were purified successfully, suggesting the ability to fold independently into stable structures. The Cas6 (C-terminal) domain retained the ability to cleave crRNA (Figure S2B) and bound the CRISPR and hairpin RNA species with higher affinity than the intact Csb2 protein (Figure 2D & 2E). In contrast, the Cas5-like N-terminal domain did not bind the CRISPR RNA, hairpin or 5’-handle species (Figure 2D, 2E, 2F). These observations strengthen the prediction that Csb2 associates with the 3’ (hairpin) end of the crRNA in the type I-G complex via interactions with the C-terminal Cas6 domain.

**Figure 2.**
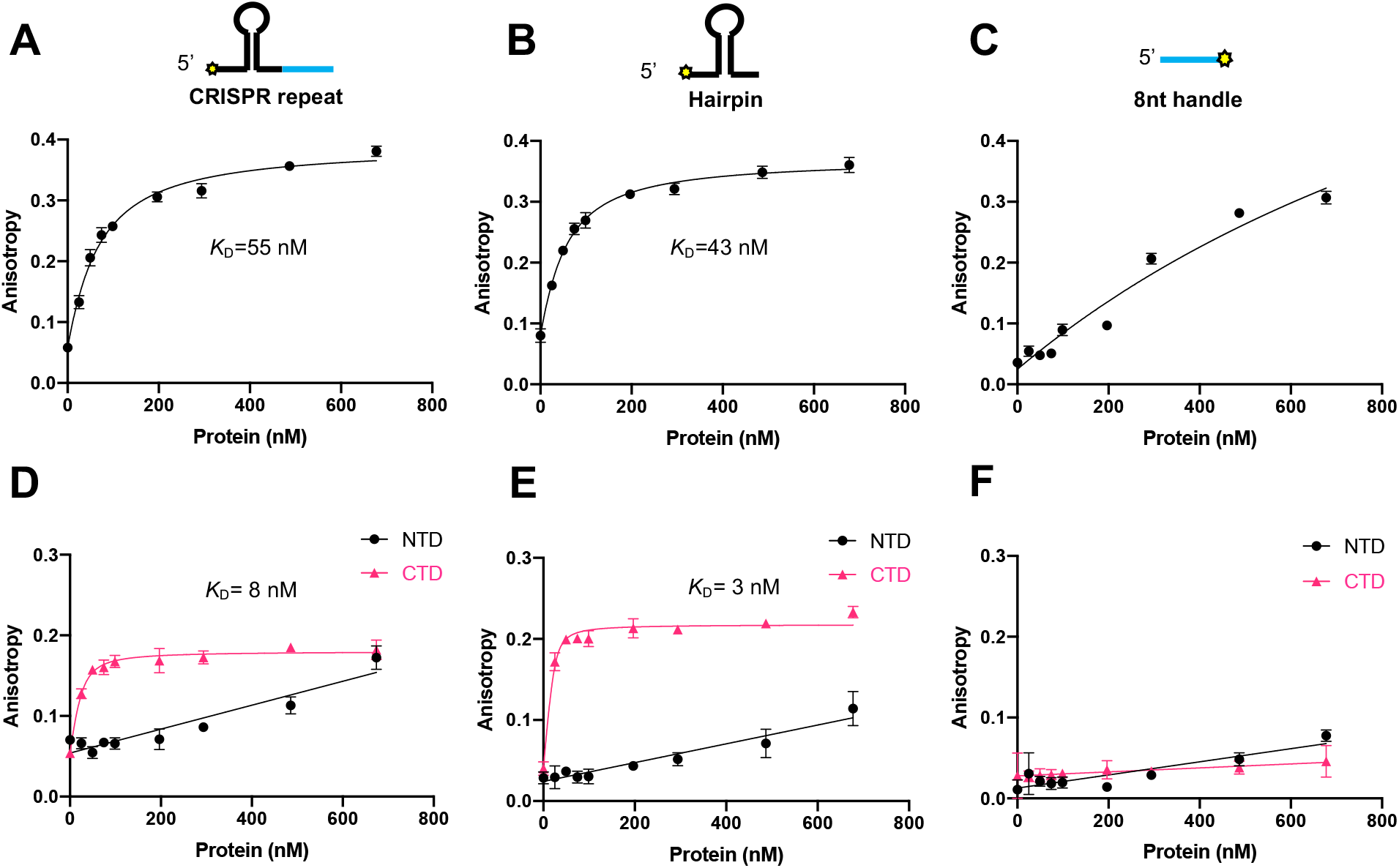
Csb2 binding affinity with CRISPR repeat, 3’ hairpin or 5’-8 nt-handle. **(A-C)** Fluorescence anisotropy analysis of Csb2 binding to the CRISPR repeat, hairpin and 5’-handle. The position of the FAM label is indicated with a yellow star. **(D-F)** Fluorescence anisotropy analysis of N- and C-terminal domain of Csb2 binding to the CRISPR repeat, hairpin and 5’-handle. Data points and error bars represent the mean of five technical replicates and standard deviation.

### Reconstruction of the effector complex *in vitro*

To determine the subunit composition and interactions of the type I-G effector complex, we incubated different compositions of subunits in the presence or absence of pre-crRNA, followed by size exclusion chromatography. We first tested the two known crRNA binding proteins, Csb2 and Cas7. In the absence of pre-crRNA, these subunits eluted separately. In the presence of pre-crRNA, Csb2 and Cas7 eluted together at an earlier time point, indicating the formation of the type I-G “backbone” in the absence of the large subunit (Figure 3A). When we repeated the experiment with Csb2, Cas7 and Cas8g, we observed the formation of a large complex “Cascade” when pre-crRNA was added (Figure 3B). Finally, we added Cas3 to the Cascade mixture, and observed the co-elution of all four subunits when crRNA was present (Figure 3C). When Cas3 was added to the backbone complex (Csb2, Cas7 and crRNA), no interaction was observed, but all four subunits co-purified when Cas8g was present (Figure S3A & S3B). Together, these experiments indicate that Csb2 and Cas7 can bind to crRNA in the absence of Cas8g, that Cas8g can bind the backbone complex to form the Cascade effector and that Cas3 is a stable component of the effector in the absence of target DNA, due to an interaction mediated by Cas8g. We also investigated the assembly and stability of the type I-G complex when Csb2 was replaced by the C-terminal Cas6 domain only. We observed no complex assembly (Figure S3C), suggesting that the N-terminal Cas5-like domain plays an important role in the formation of a stable effector complex.

**Figure 3.**
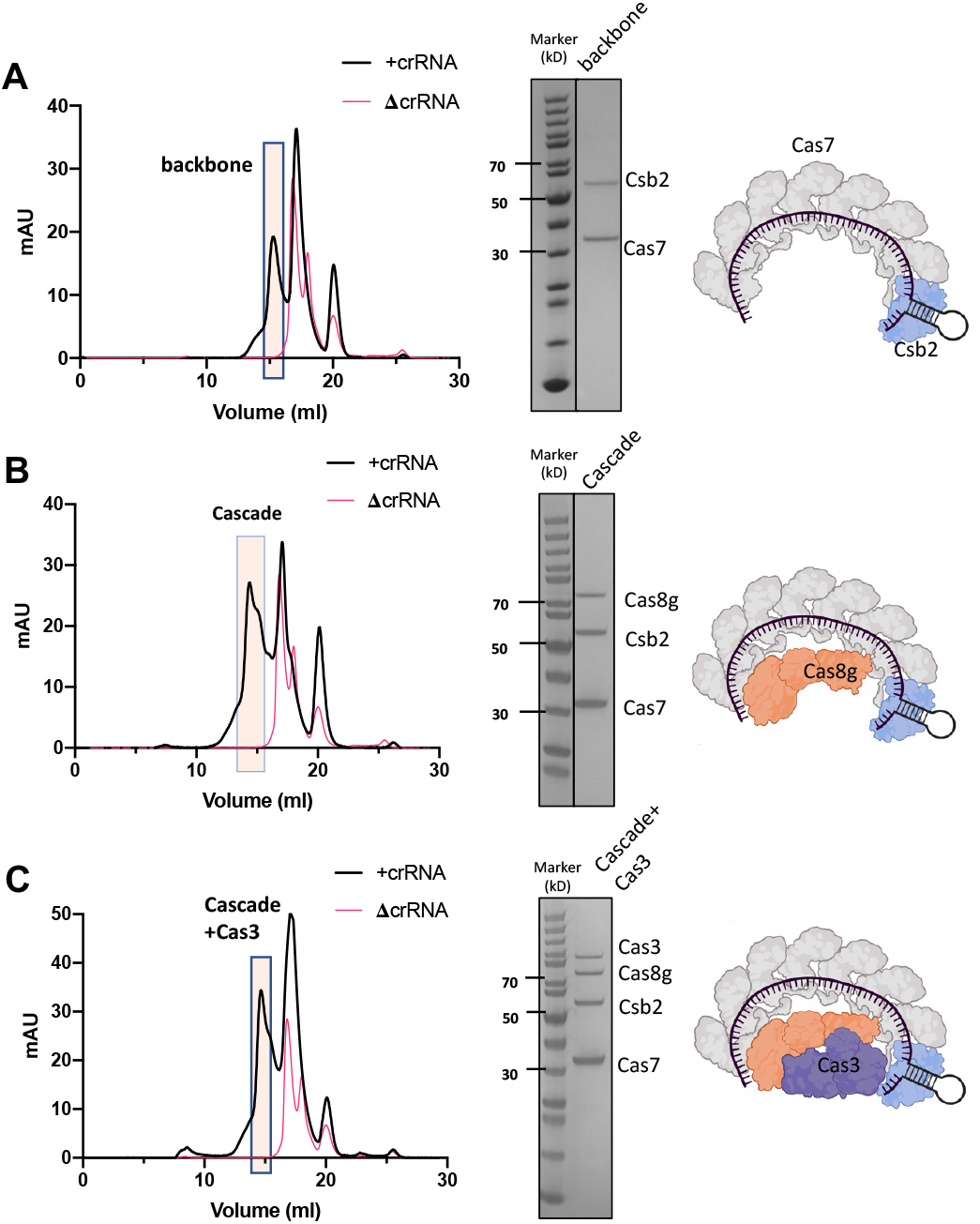
*In vitro* reconstruction of the type I-G complex. Recombinant protein subunits were incubated with *in vitro* transcribed pre-crRNA and subjected to size exclusion chromatography. Each panel shows the resulting chromatograph, SDS-PAGE analysis of the indicated fractions and schematic representation of the complex obtained. **(A)** Cas7 and Csb2 form a defined complex with crRNA. **(B)** Cas8g forms a stable Cascade complex with Cas7/Csb2/crRNA. **(C)** Cas3 forms a stable complex with the type I-G Cascade. In all cases, complex formation was dependent on the presence of crRNA.

### Target DNA binding and cleavage by the type I-G effector

With the type I-G effector reconstituted, we wished to examine the target DNA binding and cleavage properties of the effector. We first examined the cleavage of target and non-target plasmid in the presence of Cascade plus Cas3 and ATP. The target plasmid, pRAT, was incubated with increasing concentrations of the effector and analysed by agarose gel electrophoresis (Figure 4A & S4A). We observed a progressive degree of nicking of pRAT as the protein concentration increased, together with a complete loss of free supercoiled DNA, which was gel-shifted (labelled *) by the effector as well as being nicked and degraded, giving rise to a clear increase in background DNA staining (Figure 4A). In contrast, the control plasmid pCDF was only partially nicked, with little decrease in free supercoiled DNA, no gel shifting and no increase in degradation products (Figure 4A). We proceeded to investigate the role of ATP in this reaction, observing that in the absence of ATP the DNA:effector complex and the presence of nicked DNA was still observed, but the background smear of cleaved plasmid was lost (Figure 4B). These observations are consistent with the processive, ATP dependent degradation of target DNA by Cas3.

**Figure 4.**
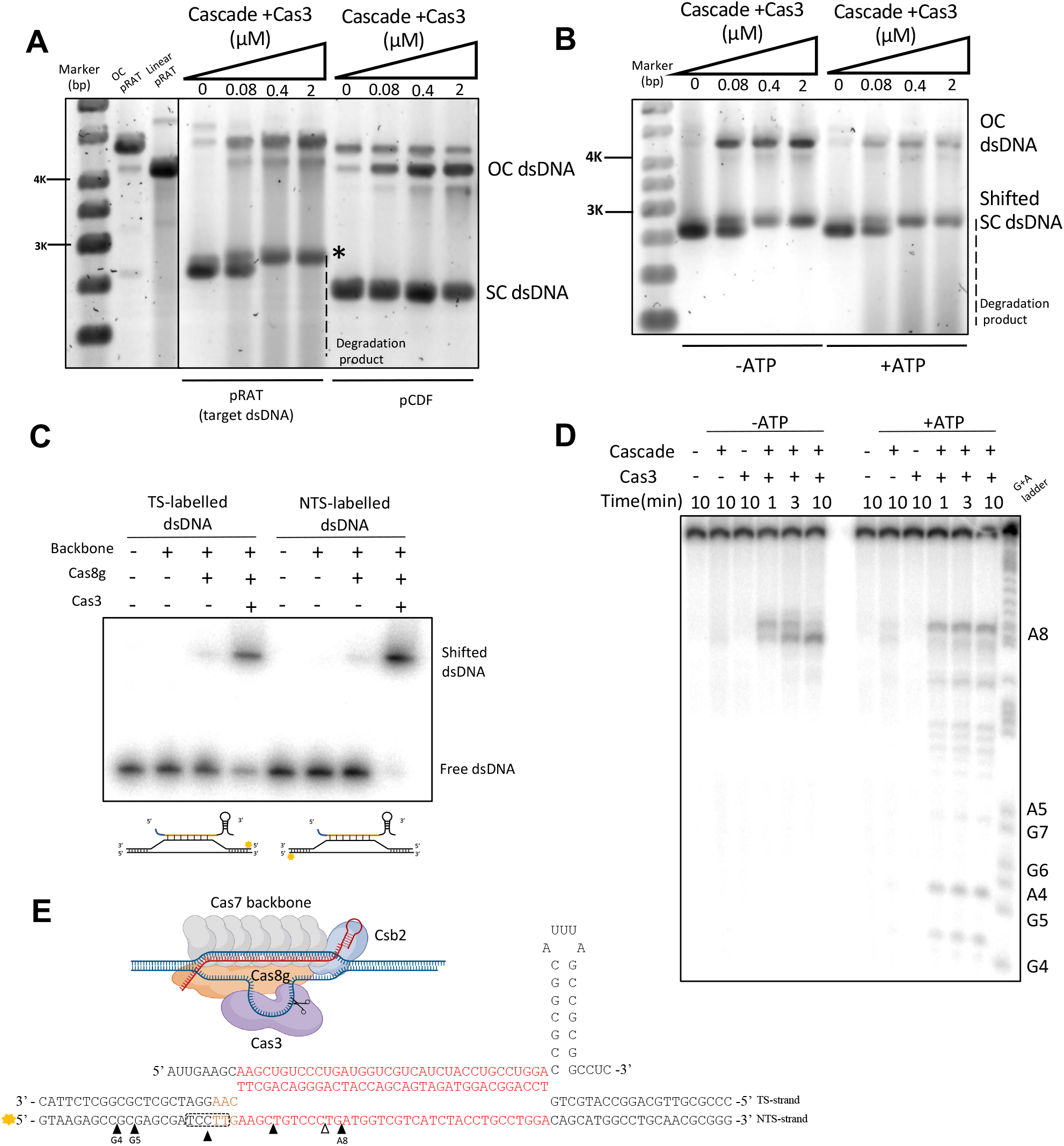
Activity of reconstituted type I-G cascade. **(A)** Supercoiled (SC) dsDNA plasmid cleavage and binding assay. Target plasmid pRAT and non-target plasmid pCDF were incubated with type I-G complex Cascade and Cas3 with ATP and analysed by gel electrophoresis. The target plasmid SC species was gel-shifted by Cascade (*), nicked to open circle (OC) form, and degraded to generate a background smear of DNA fragments. The non-target plasmid was partly nicked, but not bound or digested. pRAT obtained by nicking endonuclease digestion; linear pRAT gained by single restriction enzyme digestion. *Target SC dsDNA was bound by the effector complex. **(B)** Target SC plasmid was incubated with Cascade and Cas3 in the absence (-) or presence (+) of ATP. ATP was required for the generation of the smear of degraded DNA species. **(C)** Electrophoretic Mobility Shift Assay (EMSA) shows that type I-G Cascade only forms a stable complex with linear dsDNA targets in the presence of Cas3. **(D)** The dsDNA target was incubated with Cascade ± Cas3 in the presence and absence of ATP and analysed by denaturing gel electrophoresis. Without ATP, Cas3 cleaved the NTS in the centre of the R-loop (position A8). In the presence of ATP, Cas3 cleaves the NTS at sites 5’ of the R-loop, consistent with the 3’-5’ polarity of Cas3. **(E)** Schematic of the Cascade-target DNA complex, and mapping of cleavage sites observed for D. Black triangles show the cleavage sites when ATP is present. The five nucleotides boxed by dash lines are all cleavage sites.

We next investigated effector binding with a radiolabelled target dsDNA fragment (80 bp). In the absence of Cas8g, the backbone complex yielded no apparent shifted species, suggesting no stable complex with target DNA. When Cas8g was present, a faint shifted complex was observed. However, when Cas3 was also present the dsDNA showed near-complete gel-shifting (Figure 4C). Cas3 alone did not bind dsDNA, (Figure S4B & S4C). These data suggest that Cas3 plays a role along with Cascade in the formation of a stable complex with target DNA. Finally, we examined the cleavage of the dsDNA substrate radiolabelled on the non-target strand (Figure 4D & S4D). As expected, degradation was dependent on the presence of Cas3 and Cascade. In the absence of ATP, DNA nicking of the non-target strand was confined to a position near the centre of the R-loop (position A8). This defines the starting position for the HD nuclease. When ATP was included, degradation extended away from position A8 in a 3’ direction, indicating the unspooling of the NTS by the helicase activity of Cas3 and subsequent nicking by the HD nuclease activity (Figure 4D, E), while the TS was not cut (Figure S4D).

### In vivo reconstruction of type I-G system

The three subunits of the type I-G effector complex were cloned into the pACE vector, and the CRISPR array, targeting the tetracycline resistance gene, was constructed in pCDF, as described in the methods. These two vectors were co-transformed into *E. coli* C43 (DE3), allowing the type I-G system to be expressed and assembled *in vivo*. To detect activity against target dsDNA, a pRAT plasmid, harbouring the *cas3* gene and the tetracycline resistance gene was transformed into competent cells expressing the type I-G effector and applied on the plates with three different conditions: I. Recipients, providing the initial number of recipient cells; II. Uninduced, tetracycline in media, cells lacking tetracycline resistance cannot grow; III. Induced, subunits of type I-G system were fully induced by D-lactose and L-arabinose (Figure 5A, B). We observed a complete loss of cells that harbour Cas3 on the Transformants plate, which suggested that small amounts of type I-G effector complex due to promoter leakiness was sufficient to eradicate the target plasmid, causing loss of tetracycline resistance (Figure 5B). To exclude the possibility that toxicity of type I-G system inhibits the cell growth, a CRISPR array targeting *lacZ* was constructed in *E. coli* DH5α, which was then made competent and challenged with non-target plasmid pRAT-Duet or target plasmid pRAT-lacZ (Figure S5A). Cells transformed with target plasmid could not grow on tetracycline plates, while adequate transformants appeared on the plates for these cells challenged with non-target pRAT-Duet (Figure S5B). This plasmid challenge assay thus indicates target plasmid clearance by type I-G system.

**Figure 5.**
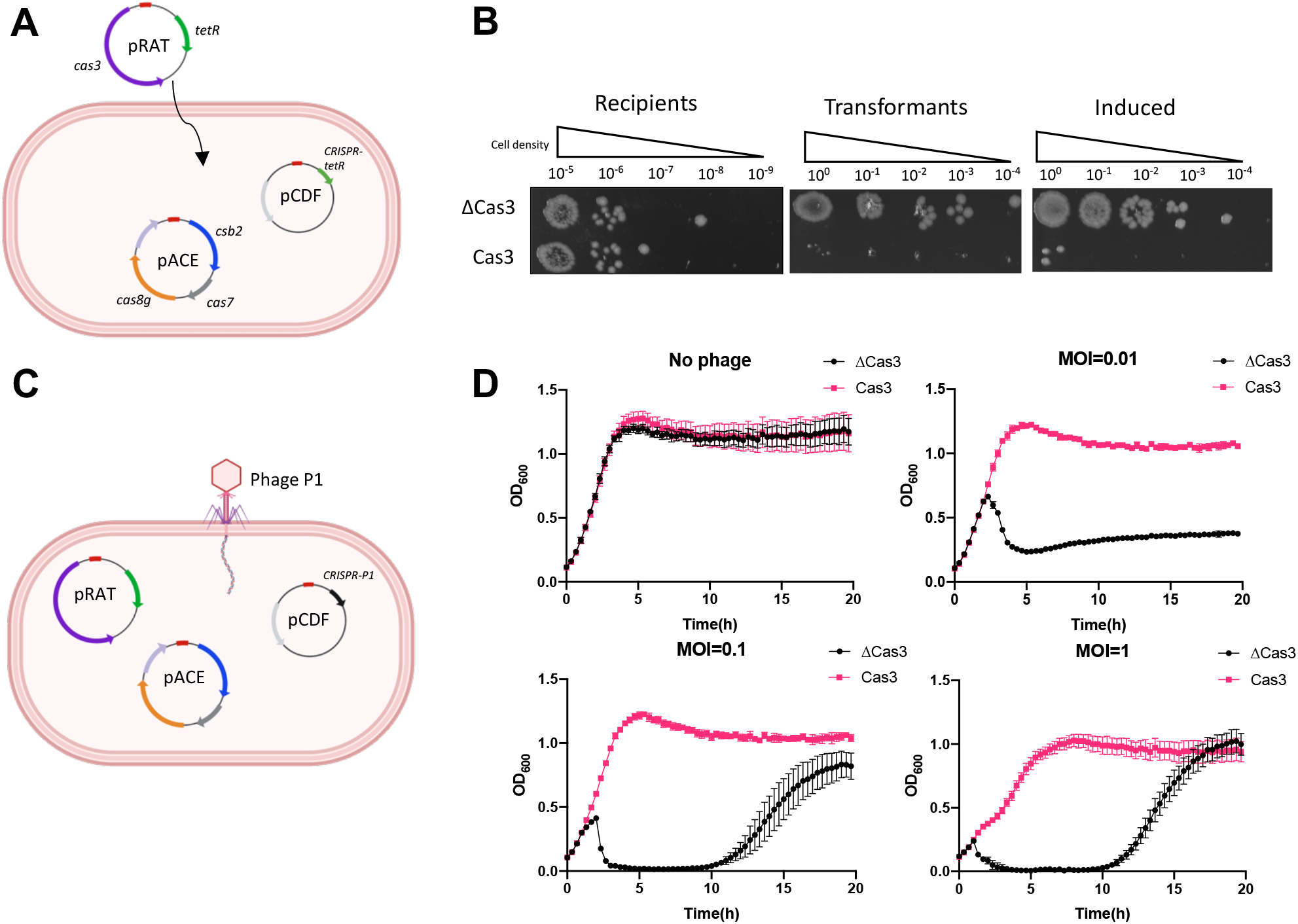
In vivo reconstruction of type I-G. **(A)** A schematic diagram explaining the plasmid challenge assay. Competent cells harbouring type I-G system were challenged with target plasmid. Csb2, Cas7 and Cas8g gene were built in pACE vector (Ampicillin resistance); CRISPR array targeting the pRAT tetracycline resistance (TetR) gene was constructed in pCDF (Spectinomycin resistance); Cas3 gene in pRAT under arabinose promoter control. **(B)** The cells on the plates in different condition. Recipients, Ampicillin and Spectinomycin in plates; Transformants, Ampicillin, Spectinomycin and tetracycline in plates; Induced, with all three antibiotics and lactose, arabinose for induction. ΔCas3, cells challenged with pRAT-Duet plasmid (Cas3 excluded). **(C)** Phage immunity assay. Type I-G system established in *E. coil*, CRISPR array targeting phage P1 *lpa* gene. **(D)** Cell growth curve, no phage infection or phage infection (MOI=0.01, 0.1 and 1). ΔCas3, cells lack of Cas3 gene. Data points represent the mean of six experimental replicates (two biological replicates and three technical replicates) with standard deviation shown.

To confirm the type I-G interference and establish the immunity in the heterologous host, we performed a phage immunity assay. Cells harbouring all subunits of type I-G complex and a CRISPR array were generated, in which the spacer targets the temperate Phage P1 late promoter activating (*lpa*) gene. Then the cells were challenged with phage P1 with three different MOI (0.01, 0.1 and 1) (Figure 5C & 5D). Across three MOI conditions, type I-G system provided immunity against phage P1, and induced type I-G system effectively inhibited the high MOI infection, fully securing cell growth (Figure 5D). On the other hand, we tested the strains in an uninduced situation. The type I-G system still protected cells from phage infection, but it only partially secured the cell growth under high load of phage infection (Figure S5C). In general, the type I-G CRISPR system was functional in the heterologous host, providing target plasmid clearance and phage immunity.

### Structural analysis of the type I-G effector complex

To understand the overall architecture of the type I-G CRISPR system and to delineate how each subunit interacts in the presence of crRNA we carried out structural reconstruction of Type I-G system using single particle Cryo electron microscopy (Figure 6 and S6). For this, the size-exclusion-purified assembled complex containing Cas7, Cas8g and Csb2 in the presence of a 72 nt crRNA was used for the structure determination. Seven interlocking Cas7 subunits which form a crescent shaped structure bound to 42 nt of the crRNA that threads through the interlocking Cas7 subunits were clearly visualised in the CryoEM density. We built the individual Cas7 subunits using the AF2 predicted structure as a starting model. The Cas7 protein adopted the typical central RAMP domain with extended β-hairpin region. Among the seven Cas7 subunits those at the 5’ and 3’ ends of the crRNA had lower resolution than the five central subunits. Cas7 subunits four and five had the highest resolution and many side chains could be visualised in the density map (Figure S7). As observed for other type I complexes, each of the Cas7 subunits occupies 6 nucleotides of the crRNA with a recurring periodic pattern of 5+1 nt, with the sixth base flipped out in the opposite direction to the remaining five. A beta hairpin emanating from the adjacent Cas7 subunit is positioned along the phosphate backbone of the sixth base, generating the kink. This structural arrangement creates an unpairing of crRNA and a target DNA at these sites. The remaining five RNA bases in each Cas7 subunit are presented for target DNA base-pairing. The crRNA is clearly visualised along the entire length of the backbone (Figure 6B). Density that may correspond to the 3’ hairpin loop is also observed, but the lower resolution precludes visualisation of nucleotides in this region. We can also trace the 5’-handle of the crRNA, which loops back and is not bound by the Cas7.1 subunit (Figure 6B).

**Figure 6.**
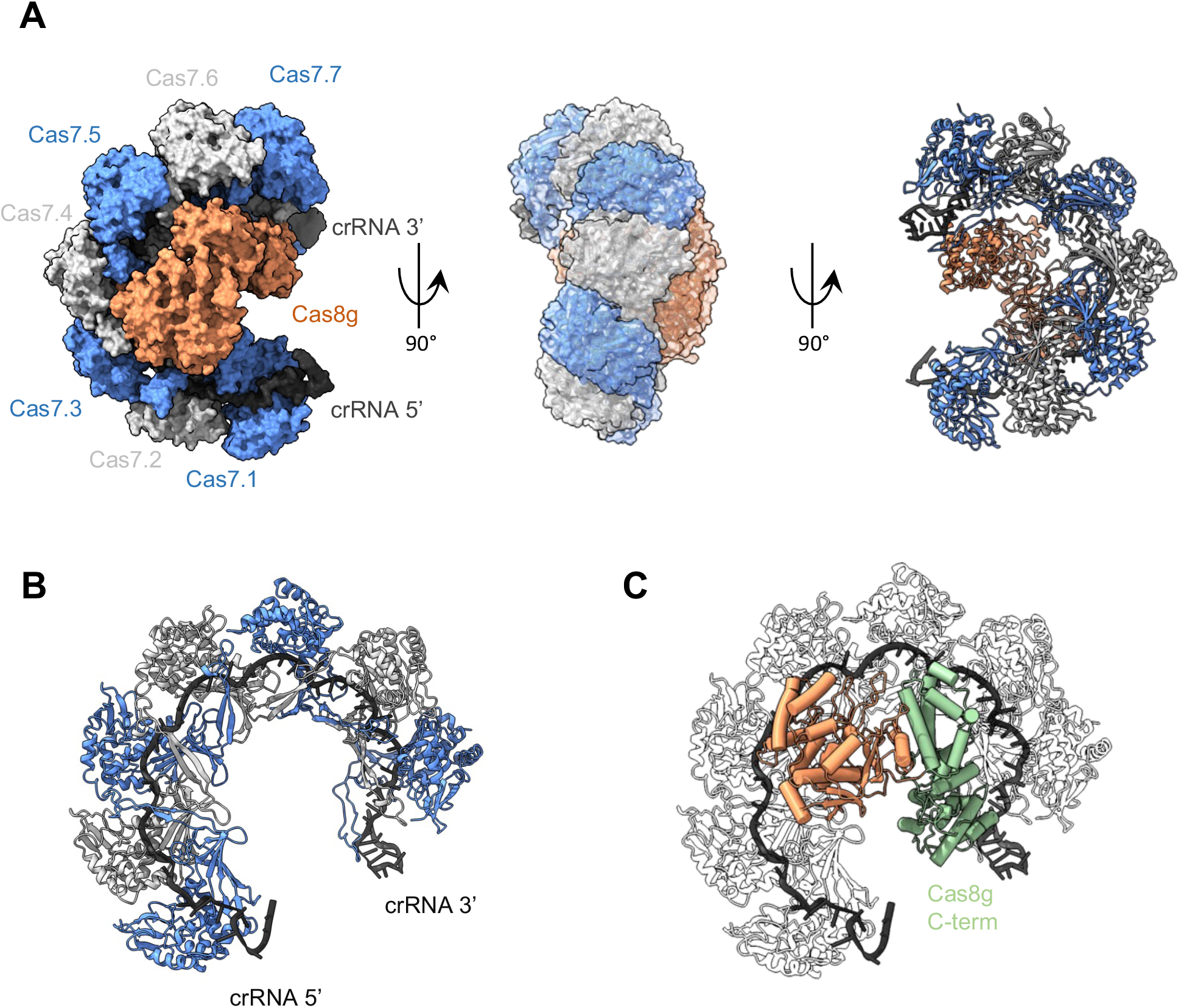
Architecture of the Type I-G Cascade complex. **(A)** 90° rotated views of the CryoEM reconstructed maps of the Type I-G Cascade showing the arrangement of seven Cas7 subunits (Blue and grey staggered), the bound crRNA (black) and the large subunit Cas8g (orange). The volume corresponding to each subunit of Cas7, crRNA and Cas8g were segmented in ChimeraX and shown in surface representation. The refined structures of Cas7, crRNA and the large subunit Cas8g were placed within the cryoEM map and shown in cartoon representation. The seven Cas7 subunits adopt a crescent shaped architecture with the large subunit Cas8g at the belly. **(B)** Refined structure of interconnected Cas7 subunits with bound crRNA shown in cartoon representation. The 5’ end of the crRNA contains 8 bases of handle which was visible closer to the edge of the Cas7.1, and the 3’ end of the crRNA constitutes 17 bases of the stem loop which was located immediately adjacent to the Cas7.7 subunit. **(C)** CryoEM refined structural model of the large subunit Cas8g shown in cartoon representation. The large subunit constitutes two distinct sub domains. The N-terminal domain of Cas8g (orange) adopts a mixed α and β fold. The N-terminal domain has extensive interaction with first two Cas7 (Cas7.1 and Cas7.2) subunits and the bound crRNA within this region. Due to lack of resolution, specific interacting residues could not be deciphered. The C-terminal domain (light green) is reminiscent of Cas11 and was placed closer to the 3’ end of the crRNA and the Cas7.7 subunit.

In our CryoEM reconstruction, we did not observe discernible density near the 3’ end of the RNA corresponding to the Csb2 subunit. As Csb2 makes up a stoichiometric component of the Cascade protein complex used for CryoEM analysis (Figure 3), we interpret the failure to observe this subunit to be a consequence of inherent flexibility of this part of the complex – as observed previously for the type I-A, I-C and I-F complexes (7,8,49).

### The large subunit Cas8g

The large subunit of type I-G systems has no detectable sequence identity to any other Cas8 protein. We predicted the structure using AF2, which gave strong support for an extended structure with a mixed (a + b) N-terminal domain and a helical C-terminal domain (Figure S8). There are no significant structural hits for this model within the protein data bank, but structural comparisons with the complete Swissprot database AF2 models using Foldseek (50) yielded a highly significant match to the predicted structure of the large Cas8a2 subunit of type I-A Cascade from *Methanocaldococcus jannaschii* and related archaea. The structural overlay shown in Figure S8 relates the Cas8a2 protein model to the N-terminal domain of the Cas8g model. The C-terminal portion is predicted to be largely α-helical, and resembles in size and secondary structure the α-helical bundle seen in the small Cas11 subunit in most Class I (type I, III and IV) effectors (51). Thus, structure predictions suggest that Cas8g is a highly divergent member of the Cas8 family with a C-terminal Cas11-like domain. This overall arrangement is also seen in the Cas8f and Cas8d proteins (15,18), and has been predicted for the Cas8g protein (52).

A poorly defined elongated density that runs across the belly of the Cas7-crRNA crescent complex structure was observed in our CryoEM reconstruction (Figure 6A). The density resembled a bi-lobed structure with a larger head closer to the 5’ crRNA end and with predominant interactions with the first two Cas7 proteins. At the 3’ crRNA end near Cas7.7, the extra density has a much thinner base. Using multibody reconstruction in RELION 4 incorporating intrinsic motion we were able to improve the overall quality of the density to ~ 8 Å resolution. The enhanced map enabled us to model the Cas8g subunit using the AF2 predicted structure as starting model. We separated the N- and C-terminal into separate rigid bodies and using DockEM we modelled the overall structure of the bound Cas8g protein. In addition, we carried out a focussed reconstruction for the Cas8g protein. For this, Cas7-crRNA core density was subtracted from the experimental particles. We then performed a 2D classification and a hierarchical 3D classification and refinement on the Cas8g isolated particles. Although the structure did not resolve to higher resolution due to relative size and the heterogeneity, the low-resolution volume showed a distinct bi-lobed structure similar to the multibody refinement structure (Figure 6C & S7G). Within different 3D classes we could observe dynamics between the two lobes. Among the three different classes two of them had an extended conformation with thinner interface between the two lobes and in one major class the lobes are closer with larger interface area. This suggests that Cas8g undergoes additional dynamic motion which could be relevant for the mechanism of Cascade.

## DISCUSSION

Type I CRISPR effectors are the most common CRISPR system present in bacterial genomes, and are increasing utilised in genome editing applications. A comparison of the published Type I CRISPR effector structures highlights the diversity of structure (Figure 7). Here, we provide the first biochemical and structural characterization of the type I-G CRISPR effector, a minimal 3 subunit system (4 including Cas3) with a unique set of features. Our work confirms the bioinformatic predictions that the Csb2 subunit is derived from a fusion between a Cas5-like and a Cas6 protein (52). The C-terminal domain of Csb2 behaves as a fairly standard Cas6 enzyme, binding tightly to the hairpin formed by the palindromic CRISPR sequence, cleaving it using a catalytic histidine residue and remaining bound to the cleaved hairpin product. The most striking divergence from the consensus is that cleavage occurs 4 nt downstream of the 7 bp stem of the hairpin rather than directly at the base. This observation should be caveated by the possibility that the crRNA adopts an extended hairpin structure when bound to Csb2. These data allow us to position the Csb2 subunit at the 3’ end of the crRNA in the Cascade complex with a good degree of confidence, despite the fact that it is not observed in the CryoEM structure. By way of comparison, type I-C Cascade uses a derived Cas5d subunit to process pre-crRNA (53), but remains bound to the 5’-handle rather than the 3’-hairpin (54). This results in the positioning of Cas5 at the 5’ end of the crRNA in the assembled type I-C Cascade, and the loss of the 3’ hairpin, presumably by hydrolysis (8). The location of a Cas5 subunit at the 5’ end of the crRNA, interacting specifically with the repeat-derived 5’-handle, is conserved in all other type I systems studied to date, so this is a major point of diversion for the type I-G effector. This divergence is exemplified by the observation that the 5’-handle of the crRNA is not directly interacting with any subunit, other than Cas7.1 (Figure 6).

**Figure 7.**
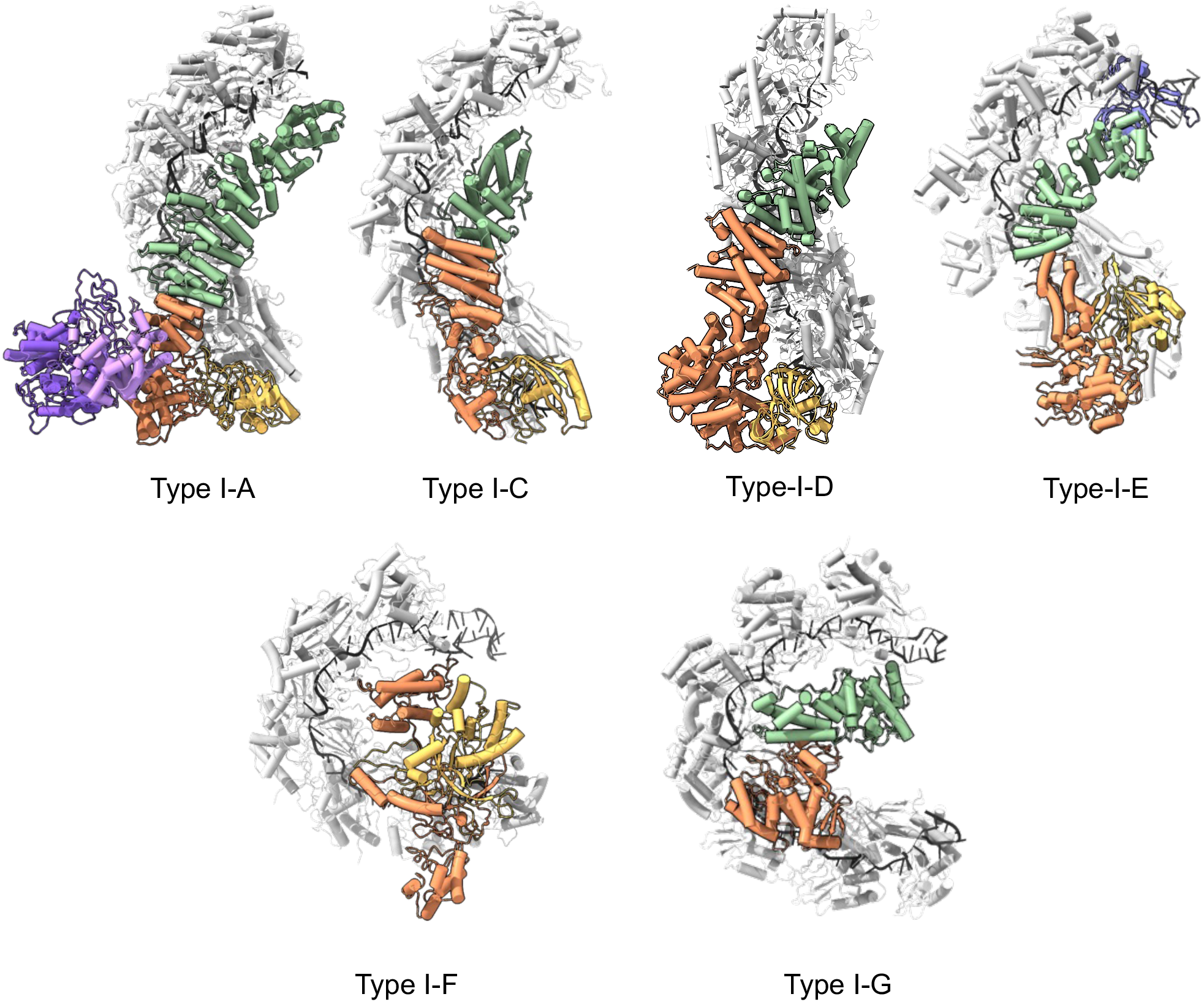
Comparison of type I CRISPR effector structures. The crRNA is shown in black and Cas7 backbone in light grey. Cas5 is shown in yellow, Cas6 in blue, Cas8 in orange, Cas11 in green. For type I-G, Cas8 N-terminal domain is orange and C-terminal domain green. For type I-A, the Cas3 HD (pink) and helicase (violet) proteins are also shown. For type I-D, Cas8 is replaced with a Cas10 subunit, also in orange, and bound target RNA is also present in the structure. Structures shown are: type I-A (7), type I-C (8), type I-D (10), type I-E (14), type I-F (49) and type I-G (this study).

With seven Cas7 backbone subunits, the type I-G effector shows pronounced curvature of the crRNA, reminiscent of that seen for type I-F systems (15) (Supplemental movie 1). Although we did not observe the Csb2 subunit, we predict that it occupies a similar position to Cas6 in the type I-E structure. The large Cas8g subunit is predicted to represent a divergent member of the Cas8 family, with detectable structural similarities to Cas8a proteins in the N-terminal portion, and a predicted Cas11-like a-helical bundle at the C-terminus that resembles the placement of Cas11 in other Cascade structures (Figure 7). The Cas8g subunit is clearly flexible and consequently is observed at only low resolution. The N-terminal domain of Cas8g sits a considerable distance from the 5’ end of the crRNA, and this together with the lack of a Cas5 subunit represents a significant point of divergence from the other structures (Figure 7). In particular, this raises a question of how PAM recognition and target DNA unwinding is accomplished by the type I-G effector. There are of course likely to be significant structural changes on target DNA binding – as observed recently for the type I-C complex (9), but there is also scope for the Cas3 subunit to play a role in these processes. As for type I-A Cascade (55,56), Cas3 is a stable component of the type I-G Cascade, likely via interaction with Cas8g in the absence of target DNA. We observed that Cas3 was required for the efficient formation of an R-loop complex between Cascade and target DNA, suggesting that Cas3 may play an active role in this process, and even perhaps in direct detection of the PAM sequence.

Our study has revealed key new information on the structure, biochemistry and unique features of the previously uncharacterised type I-G effector complex. Important questions for future studies are the placement of the Csb2 subunit (in particular, the role of the N-terminal Cas5-like domain which remains elusive) as well as the structural reorganisations and roles of the Cas8g and Cas3 subunits in target DNA recognition.

## ACCESSION NUMBERS

Atomic coordinates and structure factors for the reported crystal structures have been deposited with the Protein Data bank under accession number 8ANE, 8ANZ and the Electron Microscopy data bank under accession number EMDB-15540.

## SUPPLEMENTARY DATA

Supplementary Data are available at NAR online.

## ACKNOWLEDGEMENT

We thank Dr Sabine Grüschow and Dr Gaëlle Hogrel for advice and discussions.

## Supplementary Data

**Figure S1.**
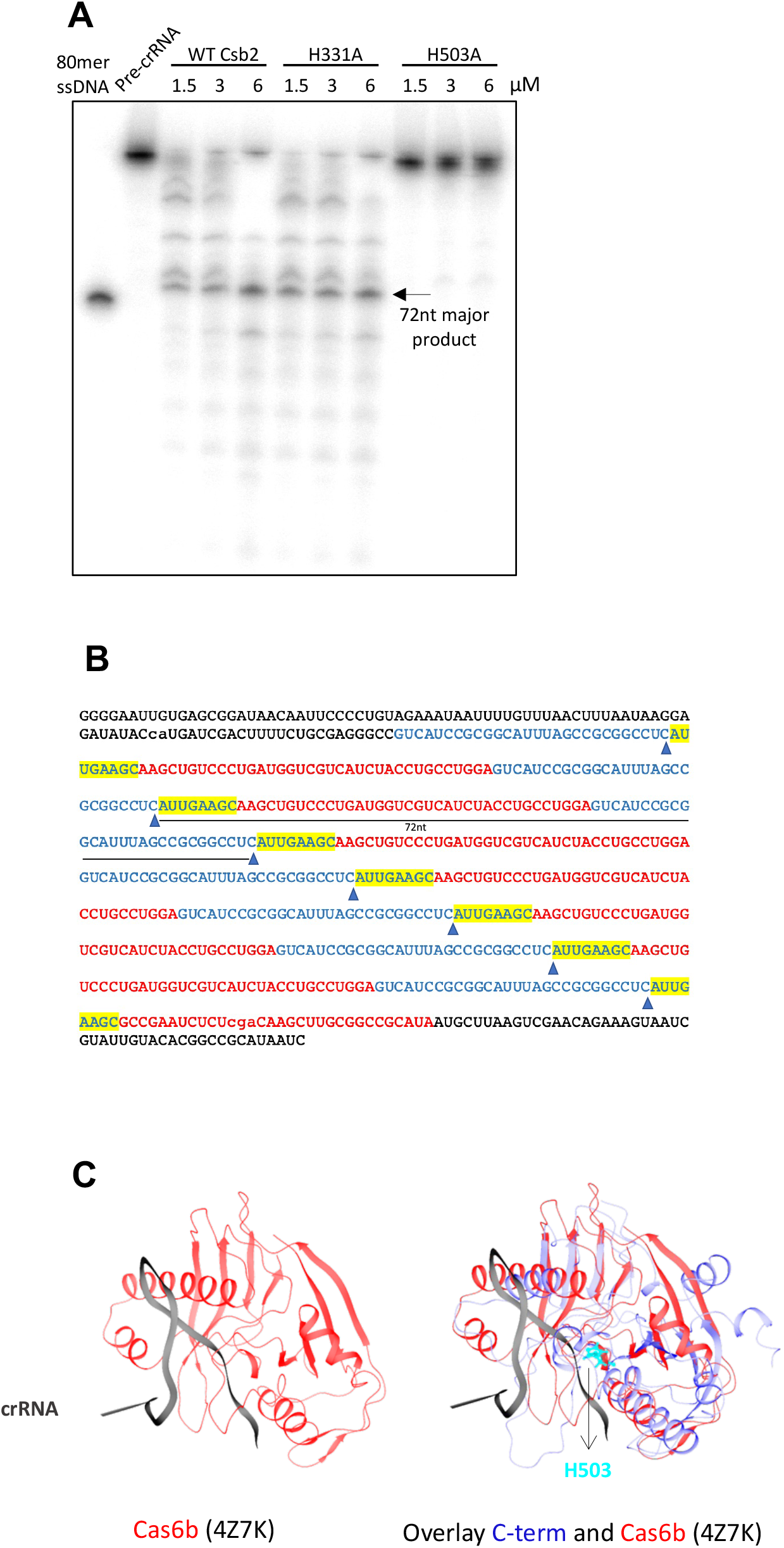
pre-crRNA cleavage by Csb2. **(A)** *In vitro* transcribed 643 nt pre-crRNA was cleaved by WT-Csb2 or mutant of Csb2. The 80 nt DNA marker (left) provides an approximate size marker for the processed crRNA. **(B)** The sequence of this 643 nt pre-crRNA, arrow indicates the cleavage site. **(C)** Structural comparison of the modelled Csb2 C-terminal domain (blue) with the Cas6b protein from *Methanococcus maripaludis* (red) reveals the likely site of crRNA hairpin binding (black, from the Cas6b structure) adjacent to the position of the H503 (cyan) active site residue.

**Figure S2.**
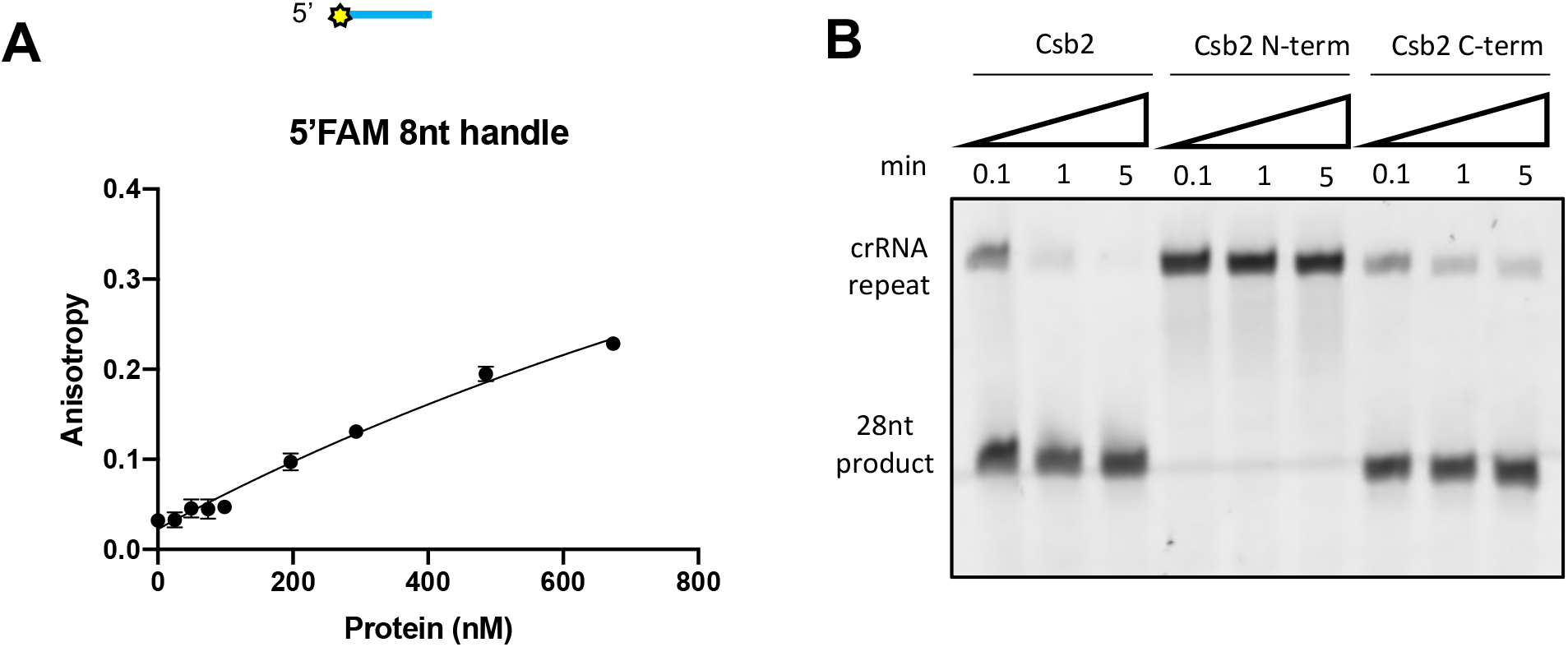
Properties of the isolated Csb2 N- and C-terminal. **(A)** Anisotropy showing Csb2 binding affinity with 5’-6FAM-labelled 8nt handle. Data points and error bars represent the mean of five technical replicates and standard deviation. **(B)** The N-terminal domain of Csb2 does not cleave the repeat, but C-terminal domain alone does.

**Figure S3.**
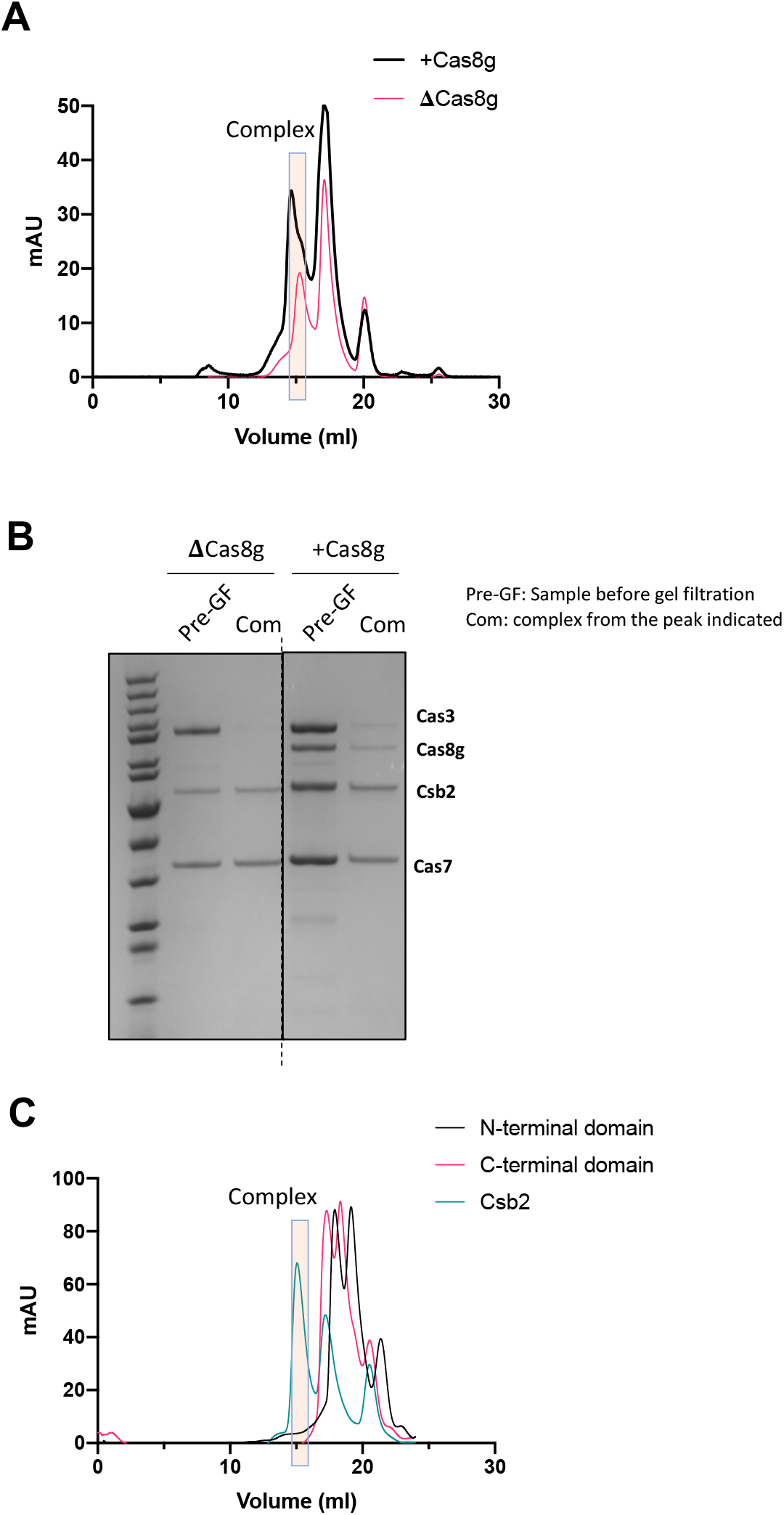
Cas3 interacts with Cas8g to be incorporated into the Cascade. **(A)** Chromatography showing complex formation in the presence of Cas8g or in the absence of Cas8g. The rectangle indicated the fraction of the complex. **(B)** The complex from chromatography was submitted to SDS-PAGE electrophoresis. **(C)** Chromatography showing only completed csb2 formed the effector complex, neither N-terminal domain nor C-terminal domain alone formed the complex.

**Figure S4.**
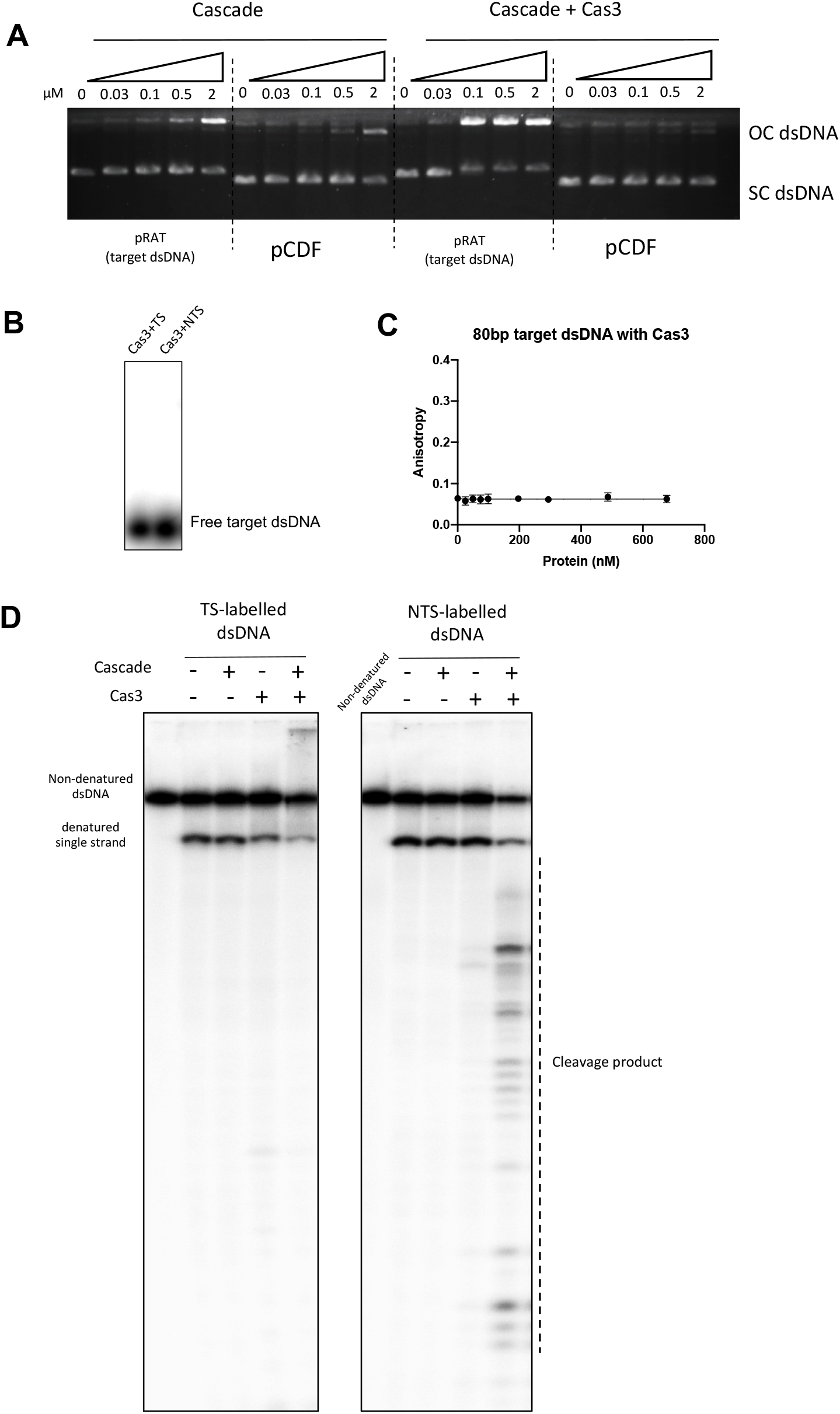
dsDNA targeting and degradation by effector complex. **(A)** Target dsDNA pRAT and non-target dsDNA pCDF were incubated with type I-G complex cascade or cascade plus Cas3, following an overnight Agarose gel electrophoresis. OC (open circular) dsDNA, SC (supercoiled) dsDNA. **(B)** Electrophoretic Mobility Shift Assay (EMS) shows that Cas3 alone has no binding with dsDNA; TS, target strand labelled dsDNA; NTS, non-target strand labelled dsDNA. **(C)** Anisotropy showing Cas3 binding affinity with target dsDNA. Cas3 alone has no binding with target dsDNA. Data points and error bars represent the mean of five technical replicates and standard deviation. **(D)** Target labelled or NTS-labelled dsDNA was incubated with Cascade and Cas3, products separated on a denaturing polyacrylamide-TBE gel.

**Figure S5.**
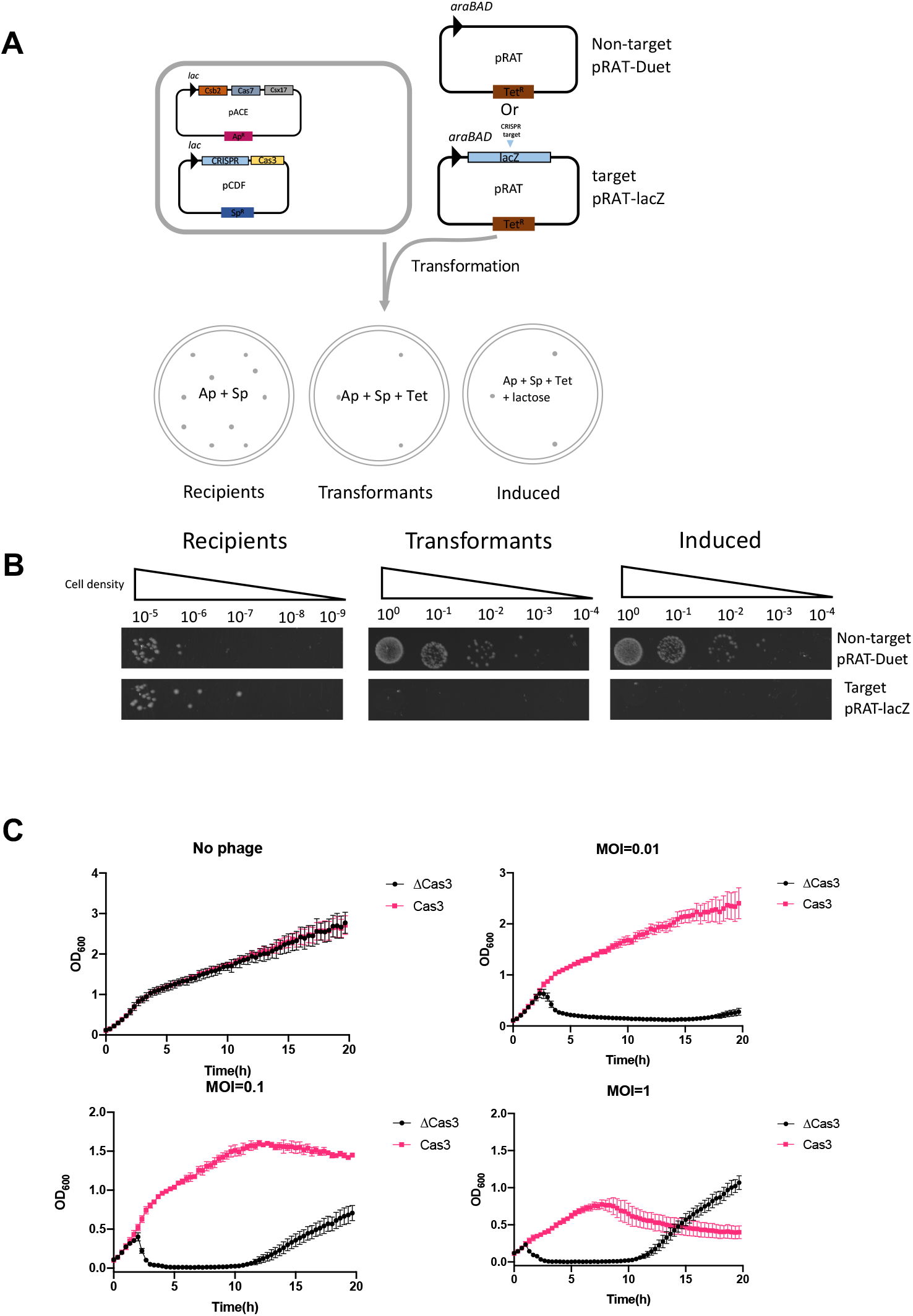
In vivo reconstruction of type I-G. **(A)** A schematic diagram showing the plasmid challenge assay for target or non-target plasmid. Csb2, Cas7 and Cas8g gene were built in pACE vector, Ampicillin resistenece (Ap); Cas3 gene and CRISPR array targeting the pRAT-lacZ were constructed in pCDF, Spectinomycin resistance (Sp); Blue triangle indicated the CRISPR array target. Recipients, Ampicillin and Spectinomycin in plates; Transformants, Ampicillin, Spectinomycin and tetracycline in plates; Induced, with all three antibiotics and lactose for induction. **(B)** The cells on the plates in different condition. **(C)** Cell growth curve without lactose and arabinose induction. No phage infection or phage infection (MOI=0.01, 0.1 and 1). ΔCas3, cells lack of Cas3 gene. Data points represent the mean of six experimental replicates (two biological replicates and three technical replicates) with standard deviation shown.

**Figure S6.**
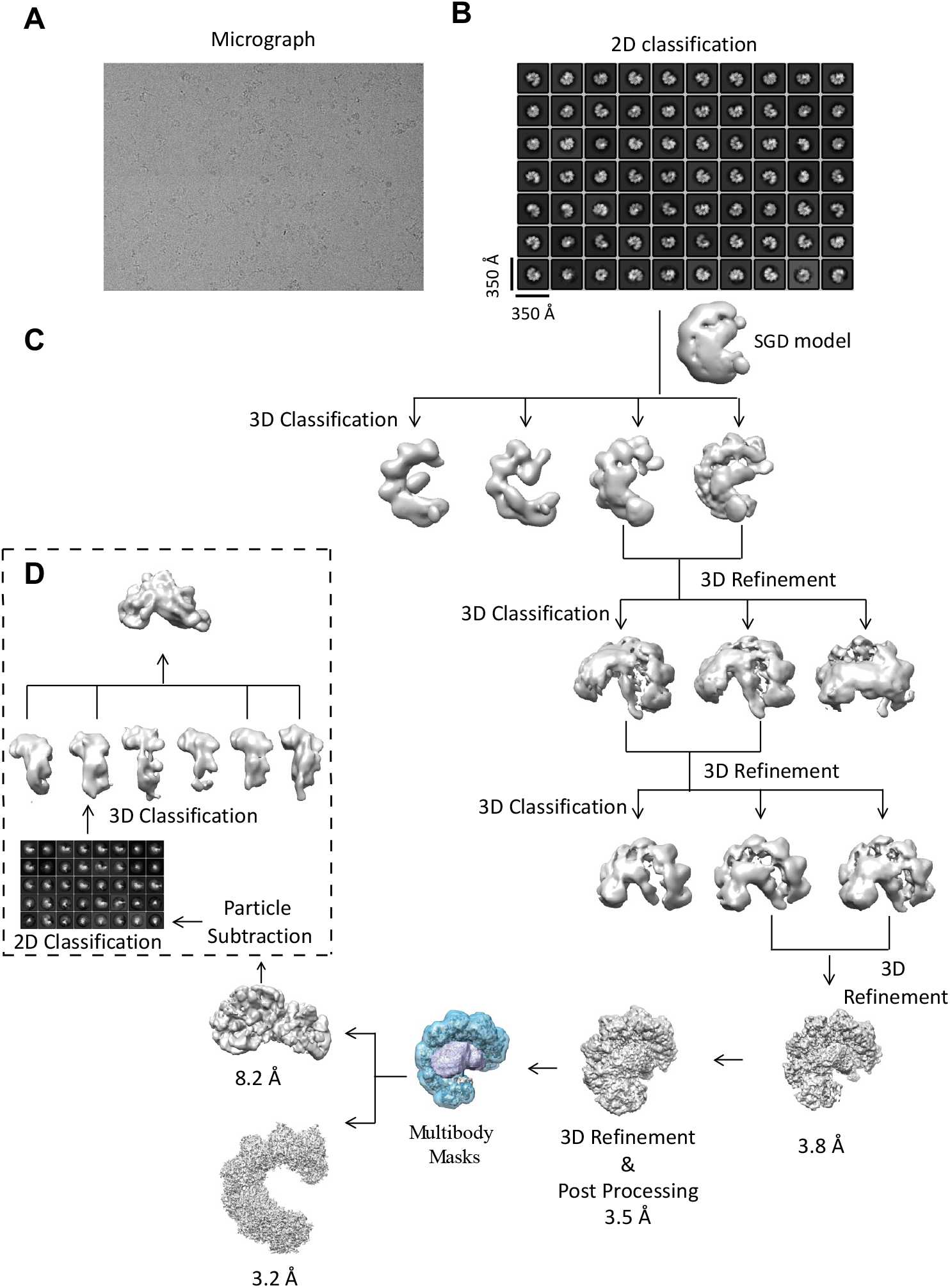
Data processing workflow for type I-G cascade reconstruction. **(A)** Representative micrograph of vitrified type I-G cascade complex collected in a Titan Krios microscope equipped with Gatan K3 detector. **(B)** Reference free 2D classification averages showing various projection images of type I-G cascade present within the dataset. A subset of these particles was used for generation of initial model using the SGD algorithm implemented in RELION 4. **(C)** Detailed workflow used to identify homogenous set of type I-G cascade complex which is refined to 3.5Å resolution. In each step, a hierarchical 3D classification was carried out. The resultant volume was then inspected in Chimera and the particles from those volumes with similar features were merged, re-extracted and a 3D refinement was carried out. At the multibody refinement step, a soft mask enclosing the Cas7-crRNA array (blue) and the large subunit Cas8g (pale green) was defined. The mask enclosed bodies were then treated as separate bodies and multibody refinement implemented in RELION 4 was carried out. The final Cas7-crRNA volume was refined to 3.2Å and the volume correspond to Cas8g was refined to 8.2Å. **(D)** Isolated particle density through particle subtraction was carried out and centred on the box (200Å) for large subunit Cas8g. Subsequently, 2D classification followed by 3D classification was carried out. 3D volumes with similar shapes were merged and a 3D auto refinement was carried.

**Figure S7.**
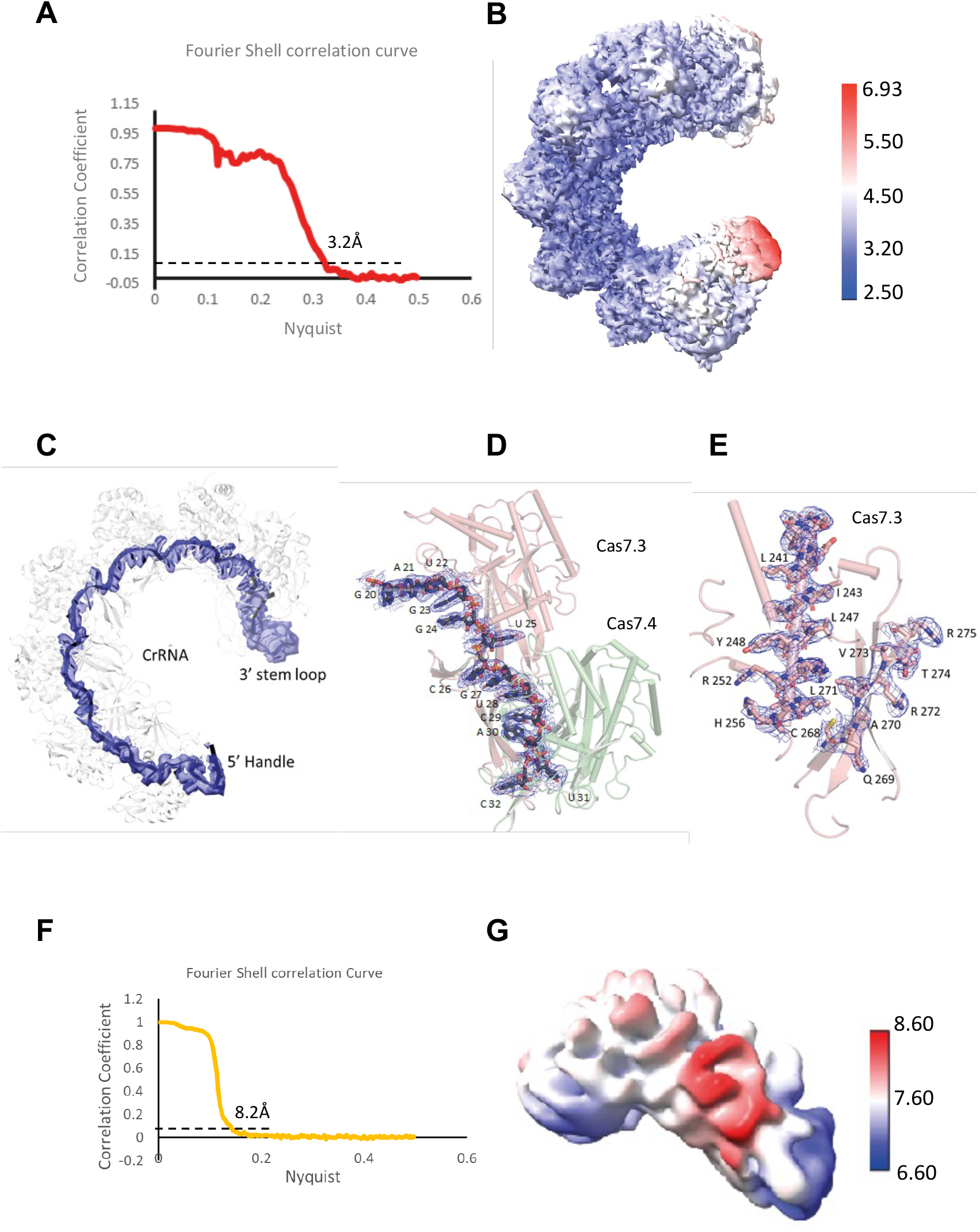
Overall quality of reconstructed Type I-G cascade complex. **(A)** Gold standard Fourier Shell Correlation curve using half maps. The estimated overall resolution for the reconstruction is 3.2A by the FSC 0.143 criterion. **(B)** Local resolution estimation for the Cas7-crRNA core using RESMAP. The Cas7-crRNA core is shown in surface representation and the surface coloured according to the estimated local resolution value. **(C)** CryoEM density for the crRNA is drawn in surface representation with the refined crRNA shown in cartoon representation. **(D)** A closer view of interaction of crRNA segment with the selected Cas7 subunits. The Cas7 subunits are shown in cartoon representation and the crRNA drawn as stick. The CryoEM density for the crRNA segment is shown in mesh. **(E)** Selected regions of Cas7 residues side chains are shown in stick and the corresponding CryoEM density drawn in mesh. Residues are numbered. **(F)** Gold standard Fourier Shell correlation curve for the Cas8g region. **(G)** Local resolution estimation for the Cas8g volume using RESMAP.

**Figure S8.**
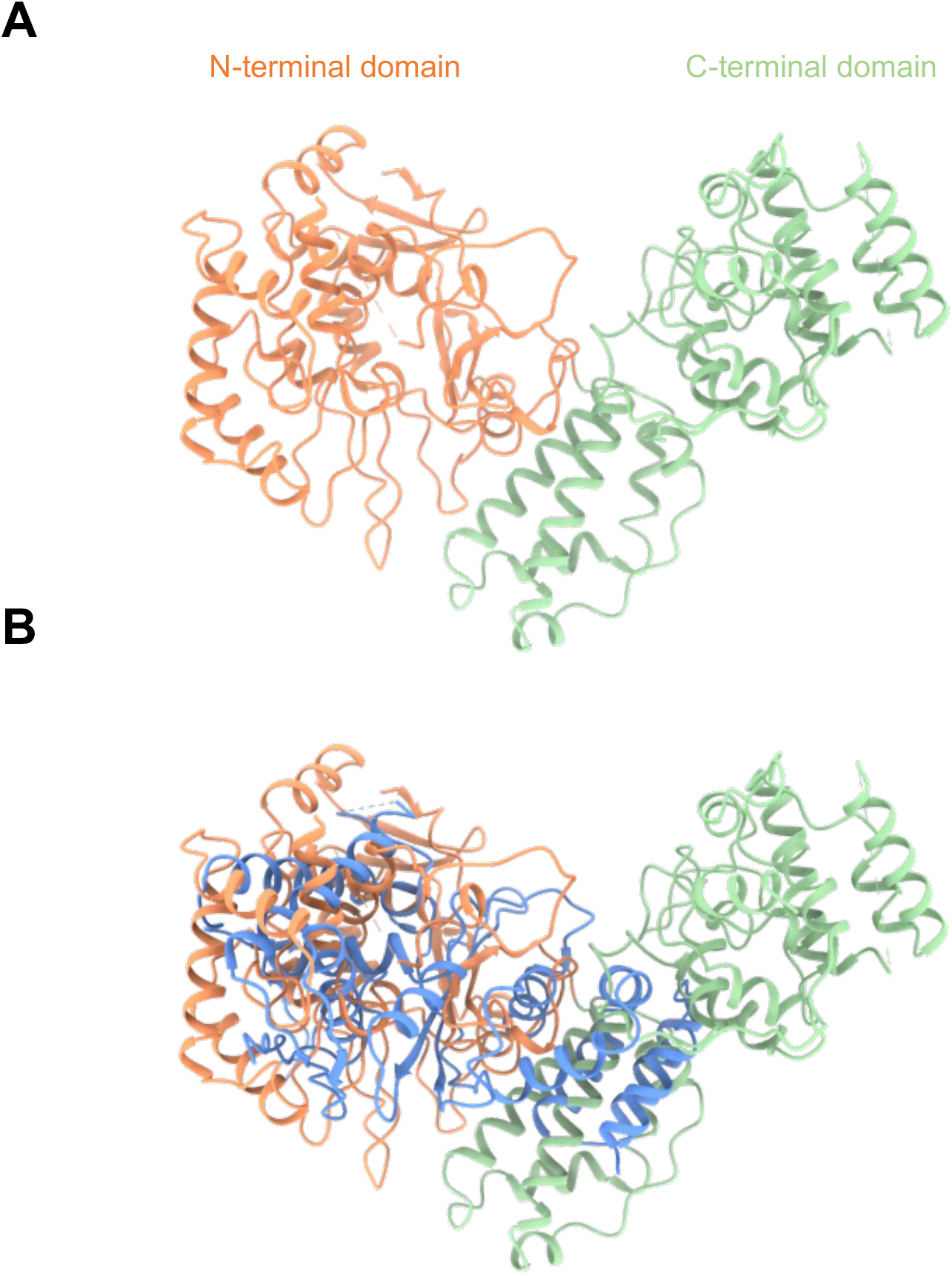
AF2 model of Cas8g/Csx17. **(A).** The N-terminal domain (orange) has a mixed a + b secondary structure while the C-terminal domain (green) is predicted for a helical bundle, similar to the composition of the Cas11 subunit in other effector complexes. **(B)** structural overlay with the AF2 model of Cas8a2 (blue) from *Methanocaldococcus jannaschii*.

**Supplementary Table 1.**
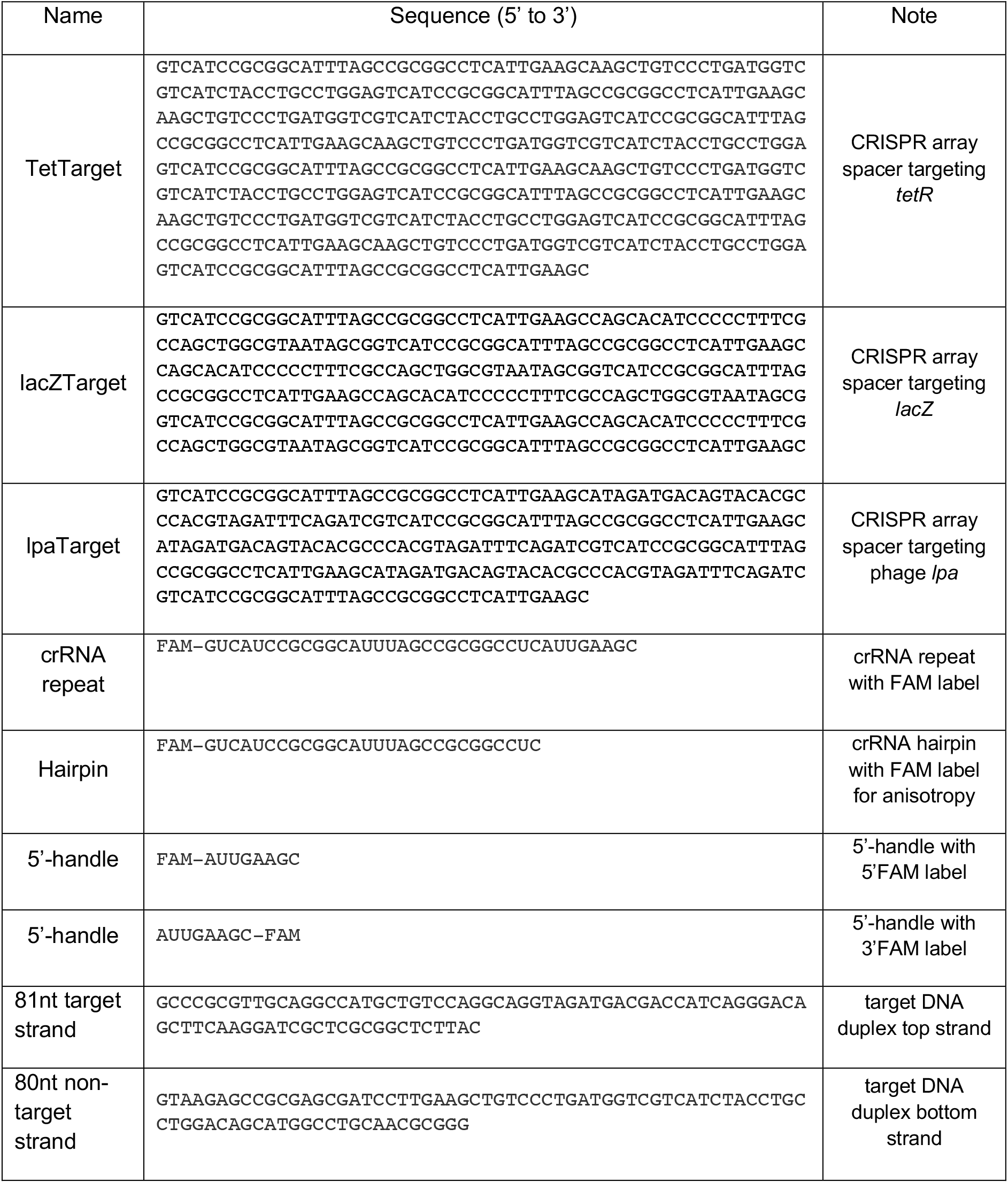
Synthetic DNA and RNA Sequences.

**Supplementary Table 2.** CryoEM Statistics.

|  |  |
| --- | --- |
| **Data collection** |  |
| Magnification | 105,000 |
| Voltage (kV) | 300 |
| Electron exposure (e-/Å<sup>2</sup>) | 40 |
| Frames in movies (no.) | 36 |
| Defocus range (µm) | 1.5-3.2 |
| Pixel size (Å) | 0.84 |
| **Reconstruction** |  |
| Symmetry imposed | C1 |
| Initial particle images (no.) | 3,420,096 |
| Final particle images (no.) | 415,000 |
| Map resolution (Å) | 3.8 |
| FSC threshold | 0.143 |
| Map resolution range (Å) | 3.8-8 |
| Sharpening B-factor (Å<sup>2</sup>) | -188 |
| **Refinement and Validation** |  |
| Subunits | Cas7, Cas8g (Csx17), Nucleotide |
| Model composition |  |
| Number of Chains | 8 |
| Protein residues | 2212 |
| RNA( no.bases) | 66 |
| Non-hydrogen atoms | 36740 |
| Hydrogen atoms | 18195 |
| **Root mean square deviation** |  |
| Bond lengths (Å) | 0.008 |
| Bond Angles (Å) | 1.025 |
| MolProbity Score | 1.23 |
| Clashscore | 3.6 |
| Poor rotamers | 0.31 |
| Ramachandran Plot |  |
| Favoured (%) | 95.2 |
| Allowed (%) | 4.8 |

**Supplementary Movie 1.**

Comparison of crRNA structures from type I CRISPR effectors, aligned at the 5’ end of the crRNA.

